# Loss of FBXO11 function establishes a stem cell program in acute myeloid leukemia through dysregulation of mitochondrial LONP1

**DOI:** 10.1101/2022.09.10.507366

**Authors:** Angela Ya-Chi Mo, Hayle Kincross, Xuan Wang, Linda Ya-Ting Chang, Gerben Duns, Franziska Mey, Zurui Zhu, Harwood Kwan, Tammy Lau, T. Roderick Docking, Jessica Tran, Shane Colborne, Se-Wing Grace Cheng, Shujun Huang, Nadia Gharaee, Elijah Willie, Jihong Jiang, Jeremy Parker, Joshua Bridgers, Davis Wood, Ramon Klein Geltink, Gregg B. Morin, Aly Karsan

## Abstract

Acute myeloid leukemia (AML) is an aggressive cancer with very poor outcomes. To identify additional drivers of leukemogenesis, we analyzed sequence data from 1,727 unique individual AML patients, which revealed mutations in ubiquitin ligase family genes in 11.2% of adult AML samples with mutual exclusivity. The Skp1/Cul1/Fbox (SCF) E3 ubiquitin ligase complex gene *FBXO11* was the most significantly downregulated gene of the SCF complex in AML. FBXO11 catalyzes K63-linked ubiquitination of a novel target, LONP1, which promotes entry into mitochondria, thereby enhancing mitochondrial respiration. Reduced mitochondrial respiration secondary to *FBXO11* depletion imparts myeloid-biased stem cell properties in primary CD34^+^ hematopoietic stem progenitor cells (HSPC). In a human xenograft model, depletion of FBXO11 cooperated with *AML1-ETO* and mutant *KRAS*^G12D^ to generate serially transplantable AML enriched for primitive cells. Our findings suggest that reduced FBXO11 primes HSPC for myeloid-biased self-renewal through attenuation of LONP1-mediated regulation of mitochondrial respiration.

## INTRODUCTION

Despite the introduction of new targeted therapies for acute myeloid leukemia (AML), outcomes remain dismal with 5-year survival being about 30% overall, and around 10% in those over 60 years old^1^. Various sequencing studies have generated molecular classification systems for AML^2,3^. Nevertheless, in a large targeted sequencing study comprising 1,540 AML patients, which established 11 distinct molecular subclasses of AML based on cytogenetic abnormalities and driver mutations, at least 11% of the patients could not be classified into a specific group^3^. Analyses of gene expression studies have suggested that additional genetic, epigenetic and/or clinical parameters play a role in refining genetic classification, and it has been suggested that half of all driver mutations in cancer remain to be discovered^2,4,5^.

Disruption of the ubiquitin pathway plays an important role in cancer development, as suggested by recurrent genetic aberrations affecting E3 ligases and deubiquitinases in multiple cancers^6–9^. Ubiquitination plays a crucial role in the development and function of normal hematopoietic stem and progenitor cells (HSPC) and leukemic stem cells (LSC), but ubiquitin gene mutations are thought to be uncommon and do not currently constitute a molecular subclass in AML^10–14^.

The presence of a ubiquitin pathway mutation has generally been extrapolated to mean that the ubiquitin-proteasome system is defective^9,15,16^, but, it is clear that different ubiquitination lysine linkages may also result in activation of protein function. Moreover, results of clinical trials with proteasome inhibitors are conflicting, and suggest that ubiquitination functions independent of proteasome-mediated degradation may play an important role in AML^17^. A clear example of a ubiquitin E3 ligase involved in myeloid malignancies that has an activating role in signaling is that of TRAF6, which mediates lysine (K)63-linked polyubiquitination^18^.

A major issue in molecular characterization of AML is that it is difficult to identify subclasses that are comprised of rare mutations in many different genes across a molecular subclass. We found recurrent somatic mutations affecting multiple components of ubiquitinating enzyme complexes in the AML Personalized Medicine Project (AML PMP) cohort^2^, and confirmed these findings in the Beat AML, TCGA AML, and TARGET AML datasets totaling 1727 unique samples^19–22^. Using an approach combining mutation analyses and gene expression changes, we identified diminished FBXO11 function – a member of the SKP-CUL-FBox (SCF) E3 ligase family – to be a key aberration imparting LSC function with myeloid bias. Loss of *FBXO11* was evidenced by truncating mutations or copy number loss, and lower mRNA and protein expression in AML compared to normal HSPC. We found FBXO11 to interact with a novel SCF^FBXO11^ target, the mitochondrial protein LONP1 in the cytosol, and through K63-linked ubiquitination promote mitochondrial entry of LONP1. Conversely *FBXO11* depletion, as seen in AML, restricted mitochondrial entry of LONP1, thereby attenuating mitochondrial respiration. Consistent with previous studies showing that reduced mitochondrial activity respiration activates a stem cell program^23–26^, we found that *FBXO11* depletion induces a myeloid-biased quiescent stem cell phenotype, and in cooperation with other mutations induces myeloid leukemia in a human xenograft model. Our results identify FBXO11 as a novel regulator of HSPC differentiation, and loss of FBXO11 contributes to leukemogenesis.

## RESULTS

### Ubiquitin pathway mutations are frequent and mutually exclusive in AML

We analyzed RNA-seq data from the AML PMP dataset^2^ to identify novel recurrent mutation classes that contribute to the initiation or maintenance of AML. In addition to single nucleotide variants (SNVs) and insertion and/or deletion mutants (indels) as previously described, we identified SNVs and indels in ubiquitin pathway genes in 11.4% (16/140) of the AML PMP cohort (Figures 1 and S1a, Table S1)^3,27^. Extending the cohort to include AML PMP, TCGA, and Beat AML datasets for a total of 1062 adult AML patients revealed ubiquitin pathway gene mutations in 11.2% of adult patients (119/1062) (Figure 1, Table S1). In contrast, only 4.5% (30/665) of pediatric AML patients had ubiquitin pathway mutations (Figure 1, Table S1). Approximately 94% (140/149) of AMLs with ubiquitin pathway system mutations only had one mutation, consistent with mutual exclusivity (Figure 1), as seen with other mutation classes with related biochemical function such as methylation and splicing^3,21,28^. Loss-of-function mutations, including truncating mutations, and copy number loss of the *FBXO11* gene and other members of the Skp1-Cul1-Fbox (SCF) E3 ubiquitin ligase complex were identified in 8.6% (91/1062) of adult AML patients (Figures 1 and S1b). Although there is evidence that deregulated ubiquitination plays a role in leukemogenesis^9,29^, these current findings implicate ubiquitin pathway mutations as being an independent and distinct molecular class in AML (Figure 1).

**Figure 1.**
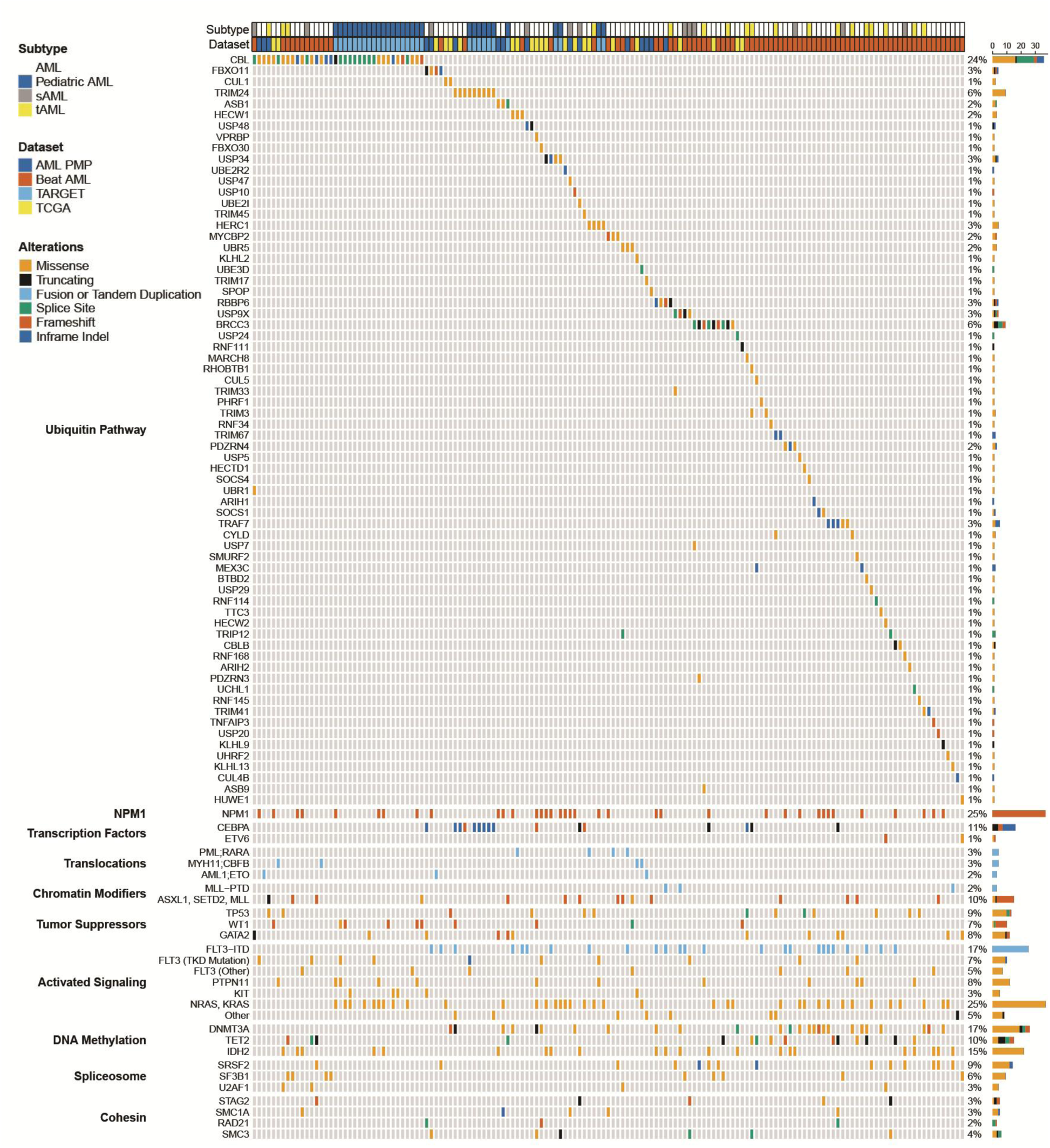
Aberration of ubiquitin pathway gene sequence or expression is frequent in AML. **a**, AML patients with ubiquitin pathway mutations from AML-PMP, TCGA, TARGET, and BEAT-AML data sets are represented in an oncoprint. Shown are 149 of 1,727 total AML patients with ubiquitin pathway mutations along with co-occurring somatic mutations, representing 11.2% of adult (AML-PMP, TCGA, BEAT-AML) and 4.5% of pediatric (TARGET) AMLs. Columns represent individual AML samples, and rows represent individual genes. Additional variants not shown are recorded in Table S1.

### *FBXO11* expression is reduced in AML

To understand potential implications of aberrant ubiquitination, we focused on the recurrently mutated *FBXO11* gene, which encodes the substrate-recognition component of the SCF^FBXO11^ E3 ubiquitin ligase complex. *FBXO11* variants occur in 1.8% of non-AML cancers in the TCGA dataset (Figure S2a, Table S2). Of the 11 patients with *FBXO11* mutations across datasets, 2 patients had a truncating or frameshift mutation and 7 patients had a single copy deletion (Figure S2b), suggesting that there is haploinsufficiency of *FBXO11* in AML. Consistent with loss-of-function mutations, AML samples with *FBXO11* mutations or deletions had lower *FBXO11* transcript expression than wild-type cases (Figure S2c).

To determine whether *FBXO11* expression was regulated in AML independent of genetic variants, we compared transcript expression of SCF genes in normal CD34^+^ HSPC and in AML cells. Of all 75 SCF genes, *FBXO11* was the most significantly downregulated gene in AML compared to normal CD34^+^ HSPC (Figure 2a, Table S3). We observed lower *FBXO11* expression across all AML subtypes (Figure 2b). Accordingly, FBXO11 protein abundance was lower in AML cell lines compared to the chronic myeloid leukemia-derived K562 cell line (Figure S2d). Importantly, bone marrow samples from AML patients had significantly reduced FBXO11 protein abundance relative to normal CD34^+^ HSPC (Figures 2c and 2d).

**Figure 2.**
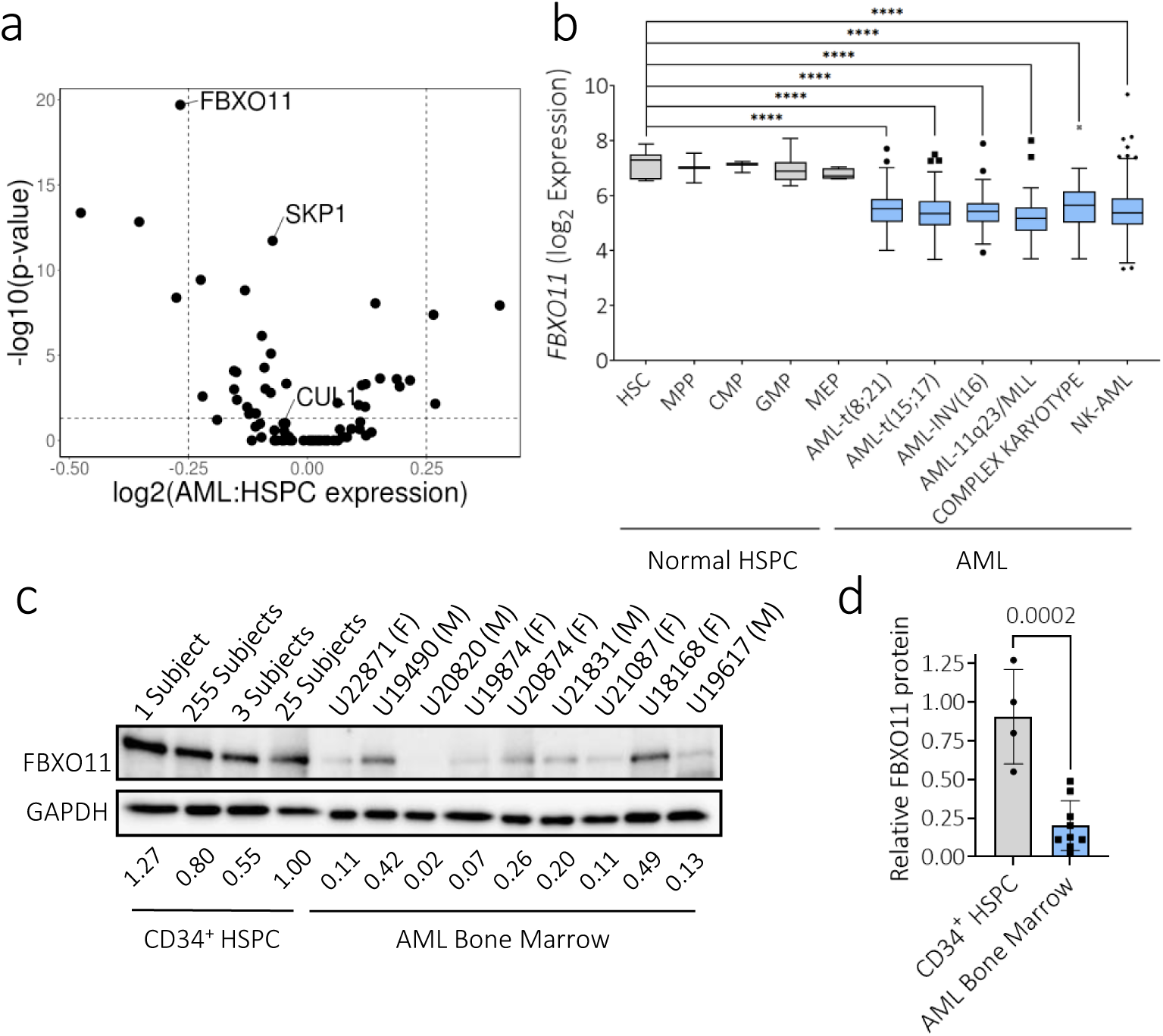
SCF gene FBXO11 expression is lost in AML compared to normal HSPC. **a**, Shown is a volcano plot comparing transcript expression of the 75 SCF genes. Bonferroni-adjusted *P* values represent *t*-tests. The horizontal dotted line represents *P* = 0.05. Vertical lines represent log_2_ fold change of |0.25|. Expression values for all SCF genes are recorded in Table S3. **b**, Expression levels of *FBXO11* transcripts across different AML subtypes and normal HSPC populations from BloodSpot are represented as a box and whisker plot. The median is shown with a horizontal line, with the edges of the box corresponding to the first and third quartiles of the distribution, with whiskers extending to 1.5* the inter-quartile range of the distribution. *P* values represent two-tailed *t*-tests, **** *P*<0.0001. **c**, FBXO11 expression in 4 normal CD34^+^ cord blood controls from 1-255 subjects and 9 AML bone marrow samples (biological sex indicated as female (F) or male (M)). Relative FBXO11 protein (fold change) is indicated normalized to GAPDH expression, and relative to the 25-subject pool. **d**, Quantification for FBXO11 expression in AML to normal CD34^+^ HSPC presented in (**c**). *P* value represent two-tailed *t*-tests, error bars represent s.d.

### Depletion of FBXO11 promotes the maintenance of primitive cells with myeloid bias

To determine the effects of *FBXO11* depletion on human hematopoiesis, we analyzed RNA expression of human cord blood-derived CD34^+^ HSPC lentivirally transduced with short-hairpin (sh)RNA targeting *FBXO11* (sh*FBXO11*) validated to deplete *FBXO11* transcript and protein, or a non-targeting shRNA (shCTR) (Figures S3a and 3b). Depletion of *FBXO11* in human CD34^+^ HSPC resulted in enrichment of HSC and LSC signatures, and diminishment of progenitor cell signatures (Figure 3b, Table S4). Interestingly, several mitochondrial-related and erythroid gene sets were also diminished compared to control (Figure 3b, Table S4). A previous study has described the requirement of FBXO11 in erythroid differentiation^30^, but not in mitochondrial function. Liquid culture of CD34^+^ HSPC resulted in increased CD34^+^ cell proportions with *FBXO11* knockdown compared to control (Figure 3c). As absolute cell numbers of CD34^+^ cells were lower with *FBXO11* depletion despite making up 60.5% ± 2% of the live cell population (Figures 3c and 3d), we postulated that *FBXO11* knockdown promoted quiescence as seen in dormant long-term stem cells. Cell cycle analysis confirmed that *FBXO11* depletion increased the G_0_ fraction in CD34^+^ HSPC (Figure 3e), which was further supported by gene signatures of quiescence (Table S4). Interestingly, although increased in the bulk cultured population of sh*FBXO11* cells, cell death was attenuated in the CD34^+^ fraction (Figure S3c), suggesting a protective effect of *FBXO11* depletion in primitive HSPC, but a cell death-promoting effect in more mature cells.

**Figure 3.**
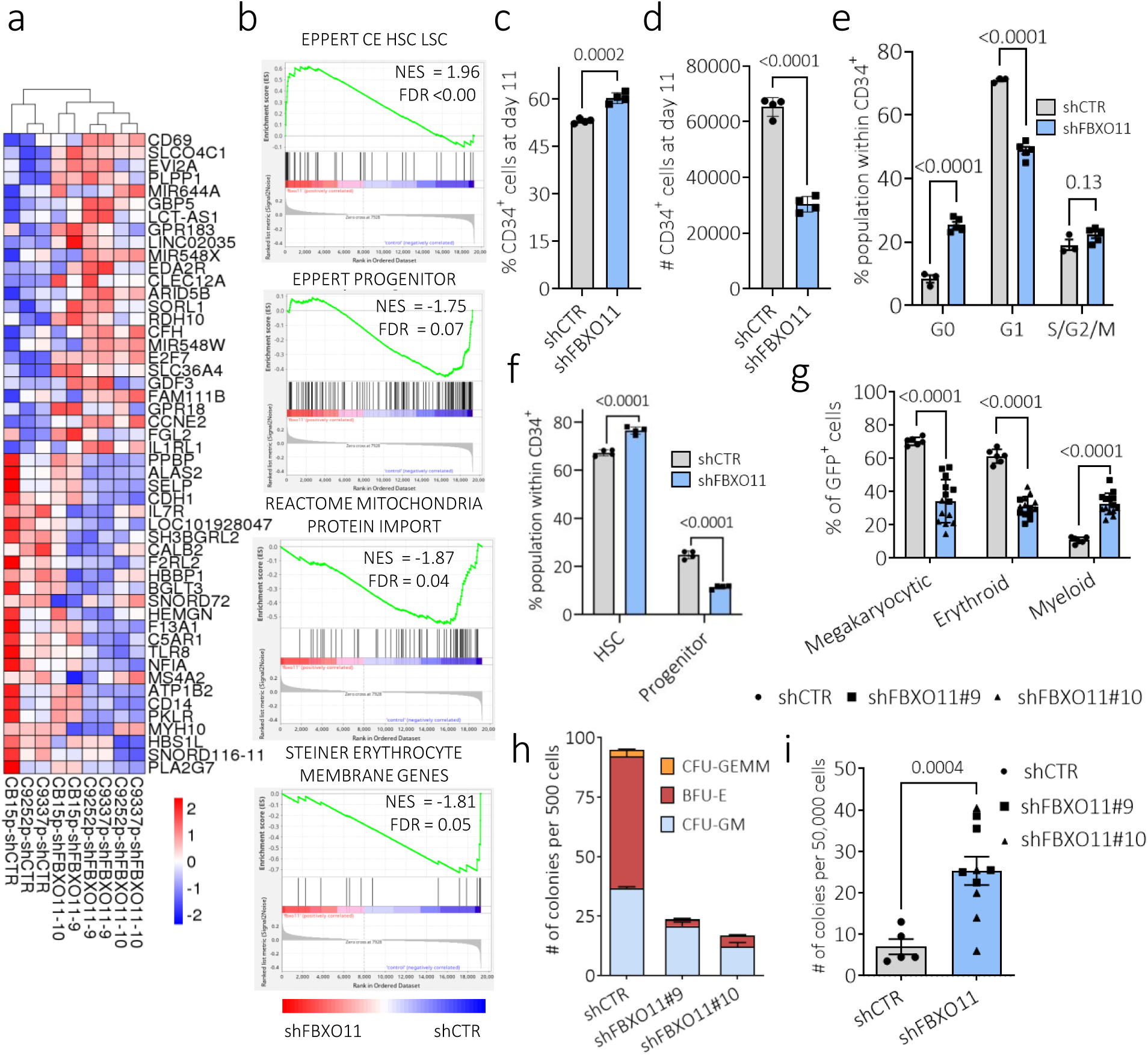
*FBXO11* depletion promotes quiescence and blocks differentiation in human HSPC. **a**, Heatmap arranged according to pairwise clustering based on the 25 most upregulated and 25 most downregulated genes in *FBXO11-*depleted cells compared to control in 3 CD34^+^ HSPC pools from independent subjects expressing a non-targeting control shRNA (shCTR) or targeting *FBXO11* (shFBXO11#9 and #10). At the bottom of the heatmap the independent HSPC pool and experimental condition are labeled. **b,** Gene Set Enrichment Analysis plots in CD34^+^ HSPC comparing *FBXO11* depletion (shFBXO11#9 and #10) to shCTR. Normalized enrichment score (NES) and false discovery rate (FDR) are indicated. All pathways passing FDR < 0.25 are recording in Table S4. **c**, Shown are percentages and (**d**) absolute cell counts of CD34^+^ HSPC expressing shCTR or shFBXO11 at day 7 of culture (*N* = 4 independent replicates). **e**, Cell cycle state of CD34^+^ HSPC was measured by Ki-67/DAPI staining (*N* = 3 (shCTR), *N* = 5 (shFBXO11) independent replicates). **f**, Percentages of CD45RA^-^ (corresponding to cultured HSC immunophenotype) and CD45RA^+^90^-^ (corresponding to cultured hematopoietic progenitor (HPC) immunophenotype) cells within the CD34^+^ population are shown (*N* = 4 (shCTR), *N* = 4 (sh*FBXO11*) independent replicates). **g**, Shown are percentages of megakaryocytic (CD41^+^ CD61^+^), erythroid (CD71^+^ Glycophorin A/B^+^), and myeloid (CD15^+^) cells in CD34^+^ HSPC expressing shCTR or shFBXO11 after 14 days of culture. **h**, Primary CFC from CD34^+^ HSPC expressing shCTR or shFBXO11 were counted 12 days after plating (*N* = 5 independent replicates). **i**, Colony counts following secondary plating of 50,000 cells from the primary CFC assay shown in panel (**f**) (*N* = 5 independent replicates). *P* values represent two-tailed *t*-test, and error bars represent s.d.

To verify a role for *FBXO11* depletion in supporting a stem cell phenotype, we examined the immunophenotype of cultured primary CD34^+^ HSPC exhibiting functional short-term and long-term stem cell (CD45RA^-^) or progenitor cell (CD90^-^ CD45RA^+^) markers^31^. *FBXO11* depletion increased the stem cell-like population, at the expense of the progenitor-like population (Figures 3f and S3d). In this culture system, all CD34^+^ cellular subpopulations showed an increase in the G_0_ fraction, consistent with *FBXO11* depletion inducing quiescence in a wide range of primitive hematopoietic cells (Figures 3e and S3e).

To examine cell output in *FBXO11*-depleted cells, we cultured CD34^+^ HSPC following knockdown of *FBXO11*. Following 14 days in liquid culture *FBXO11*-depleted cells displayed myeloid-biased output with expansion of CD15^+^ myeloid cells, but reduction in erythroid (CD71^+^/CD235a^+^) and megakaryocytic (CD41^+^/CD61^+^) populations compared to control (Figure 3g). *FBXO11* knockdown resulted in reduced primary colony forming cell (CFC) counts (Figure 3h), but increased clonogenic activity upon secondary replating (Figure 3i), consistent with activation of a quiescent HSPC population. In primary CFC assays, there was significant reduction of erythroid colonies in keeping with reduced erythroid gene expression signatures and reduced erythroid representation in liquid culture (Figures 3b and 3g, Table S4), and consistent with a known role for FBXO11 in erythropoiesis^30^. Together these results suggest that *FBXO11* depletion in human CD34^+^ cells results in a quiescent, myeloid-biased primitive hematopoietic population with stem cell characteristics.

### FBXO11 promotes mitochondrial trafficking of LONP1, a novel SCF^FBXO11^ target

To identify substrates of the SCF^FBXO11^ complex, we immunoprecipitated FLAG-tagged FBXO11 (FLAG-FBXO11) in the presence or absence of endonuclease treatment to identify proteins directly interacting with FBXO11, rather than indirectly through potential interactions with nucleic acids. We identified 5 common co-immunoprecipitating proteins using tandem mass spectrometry, including components of the SCF^FBXO11^ complex, SKP1, CUL1, and FBXO11 itself (Figures 4a and S4, Table S5). Additional proteins included the mitochondrial protease and chaperone LONP1, and the transcriptional repressor THAP5 (Figures 4a and S4, Table S5). FBXO11 interaction with LONP1 was confirmed by reciprocal co-immunoprecipitation and western blotting (Figure S5), but we were not able to confirm FBXO11 interaction with THAP5.

**Figure 4.**
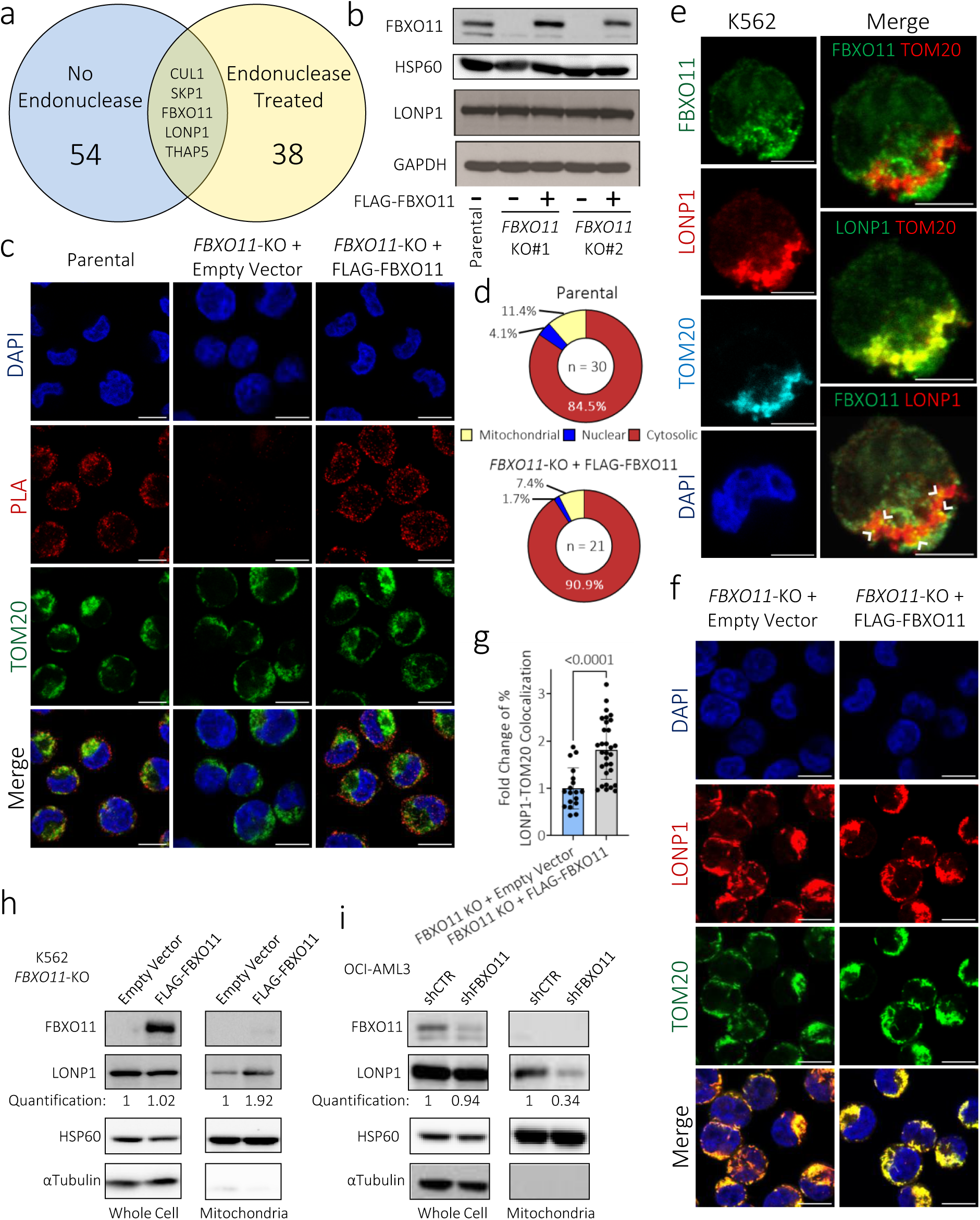
FBXO11 promotes mitochondrial trafficking of LONP1, a novel SCF^FBXO11^ target. **a**, Proteins detected by MS/MS following coimmunoprecipitation with FLAG-tagged FBXO11 (FLAG-FBXO11) in K562 cells with or without endonuclease treatment (*N* = 3 (No endonuclease) and 4 (Endonuclease treated) independent replicates). All proteins detected in each replicate are listed in Table S5. **b**, Whole cell lysates from K562 parental cells or *FBXO11*-KO CRISPR clones expressing empty vector or FLAG-FBXO11 were immunoblotted as indicated. **c**, Representative images of proximity ligation assay (PLA) for interaction of FBXO11-LONP1 in K562 parental, *FBXO11*-KO transduced with empty vector or FLAG-FBXO11, co-immunostained for TOM20 and DAPI (scale bars = 10µm). **d**, Mean percentage of nuclear, mitochondrial, and cytosolic FBXO11-LONP1 PLA in single cells represented in (**c**) from 3 independent experiments (s.d. = 2.6-5.3% (Parental) and 3.1-4.8% (*FBXO11*-KO + FLAG-FBXO11). **e**, Immunofluorescence staining of indicated endogenous proteins in K562 cells. Pairwise overlays show colocalization in yellow of indicated proteins, and FBXO11 + LONP1 colocalization is marked with arrowheads (scale bars = 5µm). **f,** Immunofluorescent staining for LONP1 localization in *FBXO11*-KO cells transduced with empty vector or FLAG-FBXO11 co-immunostained with TOM20 and DAPI (scale bars = 10µm). **g**, Fold change of percent LONP1 + TOM20 colocalization for *FBXO11*-KO + FLAG-FBXO11 relative to empty vector in single cells presented in (**f**) (*N* = 18 to 31 single cells from 3 independent experiments). *P* value represent two-tailed *t*-test, error bars represent s.d. **h,** Western blot for LONP1 expression in whole cell and isolated mitochondria lysates from *FBXO11*-KO cells transduced with empty vector or FLAG-FBXO11 or (**i**) OCI-AML3 cells expressing a non targeting control (shCTR) or *FBXO11* targeting shRNA (shFBXO11#9). Quantification represents mean LONP1 protein for FLAG-FBXO11 (**h**) (*N* = 2) or shFBXO11 (**i**) (*N* = 3, s.d = 0.06) relative to empty vector or shCTR in the indicated lysate.

To understand the impact of FBXO11 on LONP1, we generated two independent *FBXO11* knockout (KO) clones in K562 cells using CRISPR/Cas9 gene targeting. Surprisingly LONP1 protein abundance was not affected by *FBXO11*-KO, nor by reconstitution of FLAG-*FBXO11* at levels similar to the endogenous FBXO11 protein, suggesting that LONP1 is not targeted by SCF^FBXO11^ for proteasomal degradation (Figure 4b). To define the subcellular compartment of the FBXO11-LONP1 interaction, we used a proximity ligation assay (PLA), which is able to identify proteins localized within 40 nm of each other. Interaction of endogenous FBXO11 and LONP1 occurred primarily in the cytosol, external to the nucleus and the mitochondria, in parental K562 cells (Figures 4c and 4d). As expected, knockout of *FBXO11* eliminated FBXO11-LONP1 proximity, and reconstitution of FLAG-*FBXO11* recapitulated these interactions mainly in the cytosol (Figures 4c and 4d).

Given that LONP1 is recognized to be a mitochondrial protein, we further explored the subcellular localization of endogenous FBXO11 and LONP1 by confocal microscopy in parental K562 cells. FBXO11 was found to be primarily cytosolic, while the majority of LONP1 localized to the mitochondria (Figure 4e). However, there was colocalization of FBXO11 and LONP1 adjacent to the mitochondria as also suggested by the proximity ligation assays (Figures 4c and 4d). To investigate whether depletion of FBXO11 impairs mitochondrial localization of LONP1, we examined the topology of these proteins. In *FBXO11*-KO cells LONP1 protein accumulated in the cytosol, while reconstitution of FLAG-*FBXO11* resulted in a >1.8-fold increase in mitochondrial LONP1 localization (Figures 4f and 4g). We confirmed that reconstitution of FLAG-FBXO11 in *FBXO11-*KO cells does not alter whole cell LONP1 protein abundance, but specifically increases mitochondrial LONP1 by >1.9-fold, using mitochondrial fractionation and immunoblotting (Figure 4h). Importantly, there was no difference in mitochondrial FBXO11 upon reconstitution of FLAG-*FBXO11* (Figure 4h), suggesting that cytosolic interactions of FBXO11 with LONP1 result in mitochondrial relocalization of LONP1. Using another myeloid cell line, OCI-AML3, we depleted *FBXO11* by shRNA, and correspondingly observed mitochondrial-specific depletion of LONP1 compared to control (Figure 4i). Collectively these data demonstrate that FBXO11 promotes mitochondrial localization of its interacting protein LONP1.

### FBXO11 depletion reduces LONP1-mediated mitochondrial respiration in AML

LONP1 is a mitochondrial protease and chaperone protein that has been reported to target misfolded subunits of ETC complex II and complex V for degradation^32^. Consequently, *LONP1* loss reduces oxygen consumption rate and mitochondrial membrane potential (MMP)^32,33^. As we observed reduced mitochondrial signatures in *FBXO11-*depleted CD34^+^ HSPC (Table S4), we hypothesized that impaired mitochondrial import of LONP1 by *FBXO11* depletion would reduce mitochondrial respiration. Concordant with this hypothesis, significantly lower basal and maximal respiration rates were observed in *FBXO11*-KO cells compared to *FBXO 1*-*1*KO cells reconstituted with FLAG-*FBXO11* (Figures 5a-c). As expected, knockdown of *LONP1* or *FBXO11* reduced MMP in two myeloid cell lines as well as primary CD34^+^ HSPC (Figures S3b, S6b, and 5d-f). Changes in MMP were not due to changes in mitochondrial mass in *FBXO11-* or *LONP1-*depleted HSPC or *FBXO11*-KO or FLAG-*FBXO11*-reconstituted cells, as determined by MitoTracker staining and TOM20 abundance (Figures S6a and S6c). These data suggest that depletion of FBXO11 with consequent loss of mitochondria-localized LONP1 (Figures 4f-i) results in defects in the electron transport chain (ETC), which is responsible for maintaining respiration and MMP.

**Figure 5.**
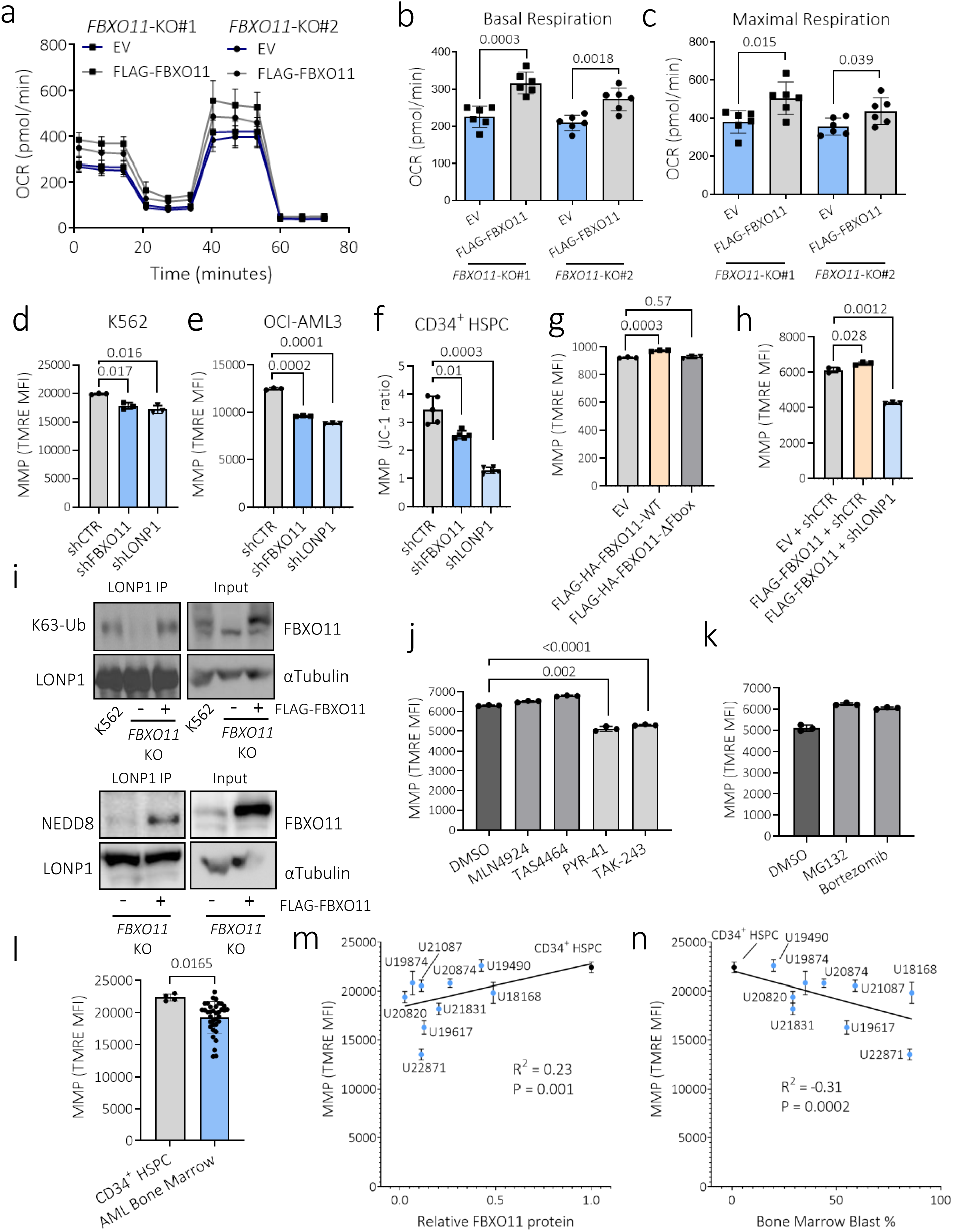
FBXO11 deficiency impairs LONP1 maintenance of mitochondrial respiration in AML. **a**, Oxygen consumption rate (OCR) measured by the Agilent Seahorse XF Cell Mito Stress Test in K562 *FBXO11*-KO cells expressing empty vector (EV) or FLAG-FBXO11. Oligomycin was added at 20 min, FCCP at 40 min, and rotenone and antimycin A at 60 min according to the manufacturer’s protocol. **b**, Quantification of basal and **c**, maximal respiration in K562 *FBXO11-*KO cells expressing EV or FLAG-FBXO11 (*N* = 6 independent replicates). **d**, Mitochondria membrane potential (MMP) measured in K562, e, OCI-AML3 cells and **f**, CD34^+^ HSPC expressing shRNA targeting *FBXO11* or *LONP1* or a non-targeting control (shCTR) (*N* = 3-5 independent replicates). **g**, MMP measured in K562 cells expressing WT *FBXO11* or *FBXO11* with deleted F-box domain (FBXO11-ΔFbox) (*N* = 3 independent replicates). **h**, MMP measured in K562 cells expressing EV with shCTR, or WT-FBXO11 co-expressed with shCTR or shLONP1 (*N* = 3 independent replicates). Immunoblots of K63-linked polyubiquitination and neddylation (NEDD8) of LONP1 immunoprecipitated from parental K562 cells or an *FBXO11* CRISPR KO clone expressing empty vector or FLAG-FBXO11 (*N* = 3 independent replicates). **j**, MMP measured in K562 cells expressing FLAG-FBXO11 treated with vehicle (DMSO), neddylation activating enzyme inhibitors (MLN4924 or TAS4464), ubiquitin activating enzyme inhibitors (PYR-41 or TAK-243), or **k,** proteasome inhibitors (MG132 or Bortezomib) (*N* = 3 independent replicates). **l**, MMP of normal CD34^+^ HSPC from the 25 subject cord blood pool and 9 AML patient bone marrow samples presented in Fig. 2c (*N* = 3-6 replicates per biological sample). *P* values represent two-tailed *t*-tests, error bars represent s.d. **m**, Correlation of sample MMP vs relative FBXO11 protein expression of the sample and **n**, Reported bone marrow blast percentage at time of clinical sampling. *P* and R values represent simple linear regression.

Long-term (LT)-HSCs and CD34^+^ HSPC with greater in vivo reconstitution potential display reduced MMP, and disruption of MMP promotes self-renewal in HSCs^26,34,24^. This is consistent with our data showing that *FBXO11* depletion promotes a quiescent stemness phenotype, which is associated with LT-HSC (Figures 3b, 3e, 3i, Table S4). In contrast, overexpression of WT FLAG-*FBXO11*, but not an F-box domain mutant (FBXO11-ΔFbox), which fails to assemble the SCF^FBXO11^ complex^35^ and disrupts interaction with LONP1 (Figure S7), increased MMP as expected (Figure 5g). To address whether LONP1 is required for the functional effects of FBXO11, we knocked down *LONP1* in cells expressing FLAG-*FBXO11*. *LON P 1*knockdown abrogated the increased MMP induced by FLAG-*FBXO11*, indicating that LONP1 is required downstream of FBXO11 to promote mitochondrial respiration (Figure 5h).

SCF^FBXO11^ is reported to act as both an E3 ubiquitin ligase as well as an E3 NEDD8 ligase ^35,36^. Given our previous data showing that LONP1 protein abundance was not affected by alteration of FBXO11 levels (Figures 4b, 4h, and 4i) and that FBXO11 activates mitochondrial LONP1 function (Figures 5a-h), we assayed for K63-linked polyubiquitination and/or neddylation of LONP1 by immunoprecipitation of LONP1 and immunoblotting for these protein-activating post-translational modifications. FLAG-*FBXO11* expression increased both K63-linked polyubiquitination and neddylation of LONP1 (Figure 5i). In order to determine whether ubiquitination or neddylation is responsible for FBXO11-driven mitochondrial function, K562 cells were lentivirally transduced to express FLAG-*FBXO11* to increase basal MMP levels, and then treated with ubiquitin E1-activating enzyme or NEDD8-activating enzyme inhibitors to determine if either mimicked the effect of *FBXO11* depletion. Only treatment with ubiquitination, but not neddylation, inhibitors reduced MMP as observed with *FBXO11* depletion (Figure 5j). As would be predicted, proteasome inhibitors also did not reduce FBXO11-mediated MMP elevation, suggesting that FBXO11-mediated, K63-ubiquitination-linked LONP1 activation, rather than proteasomal degradation, is required for regulating mitochondrial function (Figure 5k). K63-linked polyubiquitination has been shown to play a role in protein cellular localization^37^, and hence this finding is consistent with data in Figures 4f-i, showing the requirement for FBXO11 in relocating LONP1 to the mitochondria.

To determine whether mitochondrial respiration was associated with FBXO11 protein abundance in primary AML patient samples, we assayed MMP in AML marrow samples as well as normal CD34^+^ HSPC. AML samples had significantly reduced MMP compared to CD34^+^ HSPC (Figure 5l), and there was a significant correlation between FBXO11 protein abundance and MMP (Figure 5m), as well as significant negative correlation between leukemic cell burden and MMP in these samples (Figure 5n). Taken together, these data are consistent with FBXO11 ubiquitinating LONP1 through K63-linkages to maintain mitochondrial respiration. Conversely, reduced *FBXO11* as seen in primary AML cells, reduces mitochondrial respiration by attenuating mitochondrial localization of LONP1.

### FBXO11 drives mitochondrial respiration and loss of stem cell phenotype through LONP1

To confirm that FBXO11 regulates the stem cell state through LONP1, we performed RNA-seq on primary CD34^+^ HSPC expressing combinations of shRNAs targeting *FBXO11* or *LONP1* alone, or combined with enforced *LONP1* or FLAG-*FBXO11* expression (Figure S8). Samples expressing *LONP1* showed similar transcriptomic states as control shRNA or empty expression vectors (Figure 6a), suggesting that overexpression of *LONP1* is not sufficient to induce functional effects without ubiquitination by FBXO11. Similarly, samples with *FBXO11* knockdown and enforced *LONP1* expression (shFBXO11_LONP1) clustered with *FBXO11* knockdown samples (shFBXO11_9, Figure 6a), again consistent with the idea that LONP1 requires ubiquitination by FBXO11 to enter the mitochondria and function, and thus expressing LONP1 in the absence of FBXO11 does not result in overt differences in transcriptomic state.

**Figure 6.**
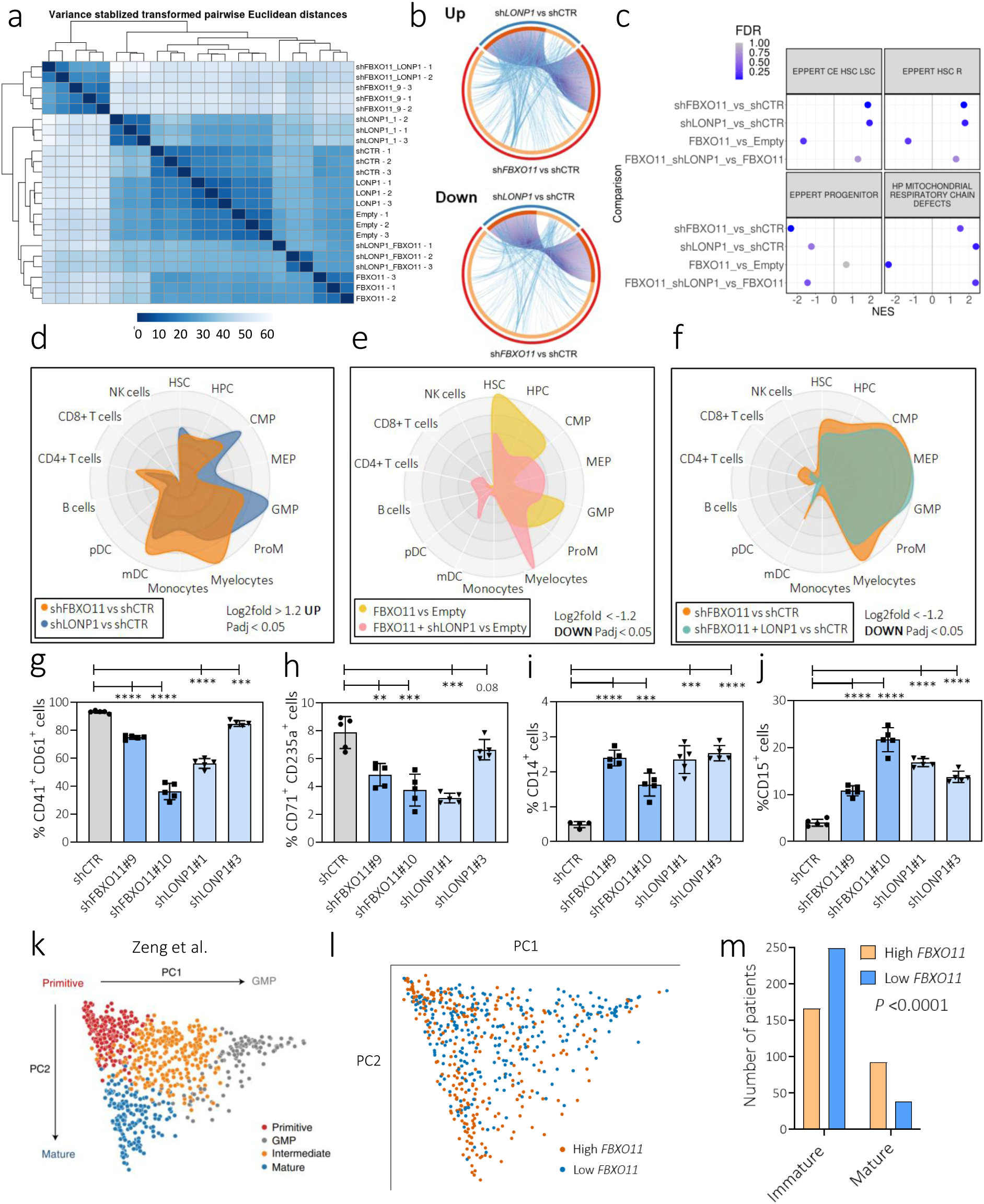
FBXO11 drives mitochondrial respiration and loss of stem cell phenotype through LONP1. RNA-seq data from CD34^+^ HSPC expressing shCTR, shFBXO11, shLONP1, empty vector (Empty), FLAG-*FBXO11*, *LONP1*, FLAG-*FBXO11* with sh*LONP1*, or *LONP1* with shFBXO11 (*N* = 3 independent replicates except for *LONP1* + shFBXO11 (*N* = 2 independent replicates)). **a**, Heatmap represents pair-wise Euclidian distances between samples based on expression of all detected genes. **b,** Circos plots comparing genes upregulated (Up) or downregulated (Down) with knockdown of *FBXO11* (red portion of outer ring) or *LONP1* (blue portion of outer ring) in CD34^+^ HSPC. Orange portions of the inner ring represent shared genes between FBXO11 and LONP1 knockdown and are connected by purple lines, yellow portions represent unique genes. Blue lines connect genes that fall under the same GO terms. Up/downregulated genes are defined by a threshold of ±1.2 log2fold change in expression, and an adjusted P value of 0.05. **c**, Enrichment trends of indicated GSEA genesets are summarized for the comparisons listed. Normalized enrichment scores (NES) for each condition are plotted. The false discovery rate (FDR) is represented by the color scale. **d-f**, Cell radar plots were generated using human normal hematopoiesis (HemaExplorer) data from the BloodSpot database compared to the significantly up or down regulated genes from the indicated conditions. The enrichment of genes associated with different hematopoietic populations indicated are compared between different groups as listed. The closer the outline is to the edge of the plot at a labeled point, the greater the proportion of differentially expressed genes associated with the corresponding cell population. (**g**) Viable CD34^+^ HSPC expressing control shRNA (shCTR), sh*FBXO11* or sh*LONP1* were assessed for megakaryocyte (CD41^+^CD61^+^), (**h**) erythroid (CD71^+^CD235a^+^) or (**i-j**) myeloid (CD14^+^ or CD15^+^) cell surface markers after 14 days of culture (*N* = 4-5 independent replicates) (*P* values represent two-tailed *t*-tests, error bars represent s.d. ** *P*<0.01, *** *P*<0.001, **** *P*<0.0001) **k**, Principle component analysis (PCA) of deconvoluted 864 de novo AML patient cellular hierarchies from Zeng et al. **l**, The 864 AML patient deconvoluted patient samples were stratified by their *FBXO11* expression into tertiles within their datasets (TCGA, BEAT-AML waves 1+2, and Leucegene) and projected by PCA, with the intermediate tertile subsequently removed. **m**, Indicated cellular hierarchy cluster for samples in Zeng et al. stratified as *FBXO11* high vs low tertiles (Immature = Primitive + Intermediate + GMP hierarchies). *P* value represents chi-square test.

Given the finding that LONP1 requires FBXO11 for its function, we compared differentially expressed genes for shFBXO11 and shLONP1. We found that 63.3% of the upregulated genes (309/488) and 52.4% of the downregulated genes (253/483) in *LONP1*-knockdown CD34^+^ HSPC were shared with *FBXO11* knockdown HSPC (Figure 6b). However, these transcripts represented only 21% (309/1470) of the significantly up- and 19.8% (253/1275) of down-regulated genes seen with *FBXO11* knockdown CD34^+^ HSPC (Figure 6b). These findings are compatible with the expectation that FBXO11 ubiquitinates multiple proteins, while LONP1 activity is dependent on FBXO11-mediated ubiquitination.

We next performed Gene Set Enrichment Analysis (GSEA) to confirm that the stem cell phenotype and mitochondrial functional effects induced by *FBXO11* depletion are dependent on LONP1 function. Both *FBXO11* and *LONP1* knockdown positively enriched for HSC/LSC gene sets as well as mitochondrial respiratory chain defects (Figure 6c). FLAG-*FBXO11* expression had the opposite effect with negative enrichment of these gene sets. However, *LONP1* depletion in FLAG-*FBXO11*-expressing CD34^+^ HSPC reversed the transcriptional effect of FLAG-*FBXO11* expression, indicating that FBXO11-directed loss of stemness is mediated by LONP1. An inverse of these transcriptional trends can be seen for the hematopoietic progenitor gene set, as noted previously (Figures 3b and 3f, Table S4). These results further support the role of LONP1 as a mediator of FBXO11-driven mitochondrial respiration and loss of stem cell phenotype.

Next, we examined whether *FBXO11* or *LONP1* knockdown enriched for a stem cell transcriptomic state and myeloid bias as seen in Figure 3. As shown in the cell radar plot, genes upregulated with knockdown of *FBXO11* or *LONP1* shared similar transcriptomic profiles, with enrichment of genes associated with HSC and myeloid cell populations including common myeloid progenitors (CMP), granulocyte-monocyte progenitors (GMP), myelocytes, and myeloid dendritic cells (mDC) (Figure 6d). Aligning with these trends, genes downregulated with FLAG-*FBXO11* expression were strongly enriched in HSC, CMP, and GMP associated genes, but *LONP1* knockdown attenuated this effect, and resulted in enrichment of the megakaryocyte-erythroid progenitor state, as expected (Figure 6e). As noted, the predicted cell states with *FBXO11* depletion combined with *LONP1* overexpression highly resembled that of *FBXO11* depletion alone, supporting a model of FBXO11-driven mitochondrial respiration that is dependent on LONP1 (Figure 6f).

To confirm the transcriptomic trends observed with *FBXO11* and *LONP1* depletion in a cellular model, we assayed differentiation of CD34^+^ HSPC in culture. Indeed, *LONP1* depletion had a similar effect as *FBXO11* depletion on primary CD34^+^ HSPC differentiation, causing a reduction in the CD41^+^CD61^+^ megakaryocytic and CD71^+^CD235a^+^ erythroid populations, and an increase in the CD14^+^ and CD15^+^ myeloid populations (Figures 6g-j), corroborating our transcriptional findings.

To determine whether *FBXO11* expression was associated with differentiation in AML patient samples, we leveraged a framework for characterizing cellular hierarchies in primary AML samples based on bulk RNA-seq^38^. Briefly, gene expression was used to deconvolute the cellular hierarchy of the sample using single cell RNA-seq references for leukemic and non-leukemic hematopoietic populations, which separates AML samples into 4 clusters based on their position in the cellular hierarchy (Figure 6k). We applied these methods to the 864 samples originally used^38^ from the TCGA, BEAT-AML, and Leucegene cohorts^19,21,39^, and stratified the samples into tertiles based on their *FBXO11* expression (Figure 6l). While individual mutations to single genes were unable to cluster samples within these hierarchies^38^, we found that the *FBXO11-*low AML samples were enriched in the 3 clusters with high proportions of immature signatures (Primitive + Intermediate + GMP) at the expense of samples in the mature cluster (Figure 6m). Collectively these findings suggest that *FBXO11* depletion induces an immature myeloid-biased state in both normal CD34^+^ HSPC and in AML cells.

### *FBXO11* depletion cooperates with *AML1-ETO* and *KRAS*^G12D^ to generate human myeloid leukemia

To determine whether *FBXO11* deficiency cooperates with driver mutations to initiate AML in vivo, we revisited our analyses of AML samples. One patient with an *FBXO11* mutation had a cooccurring *AML1-ETO* (*RUNX1-RUNX1T1*) fusion (Figure 1), and ubiquitin pathway mutations cooccur with *RAS* mutations (Figure S9). Previous studies have shown that the combination of canonical full-length *AML1-ETO* with mutant *RAS* is not sufficient to transform human CD34^+^ cells, likely due to a lack of sufficient self-renewal activity. As there is no known combination of mutants with *AML1-ETO* that results in human leukemia^40^, we asked whether the combination of *AML1-ETO*, activated *KRAS*, and *FBXO11* depletion could initiate human leukemia.

We transduced CD34^+^ HSPC with combinations of these variants: empty vector (EV), *FBXO11* shRNA (F), AML1-ETO (A), F+A (FA), *KRAS*^G12D^ (K), F+K (FK), K+A (KA), or all three (FKA) (Figures 7a and S10C). We then flow-sorted for the co-expressed fluorescent markers and placed the cells into long-term culture on MS5 cells (Figure 7a). Following long-term co-culture, *FBXO11* knockdown alone resulted in reduced cell numbers (Figure S10a), but significant enrichment of the proportion of CD34^+^ cells when combined with A or KA relative to corresponding controls (Figures 7b, 7c, and S10a).

**Figure 7.**
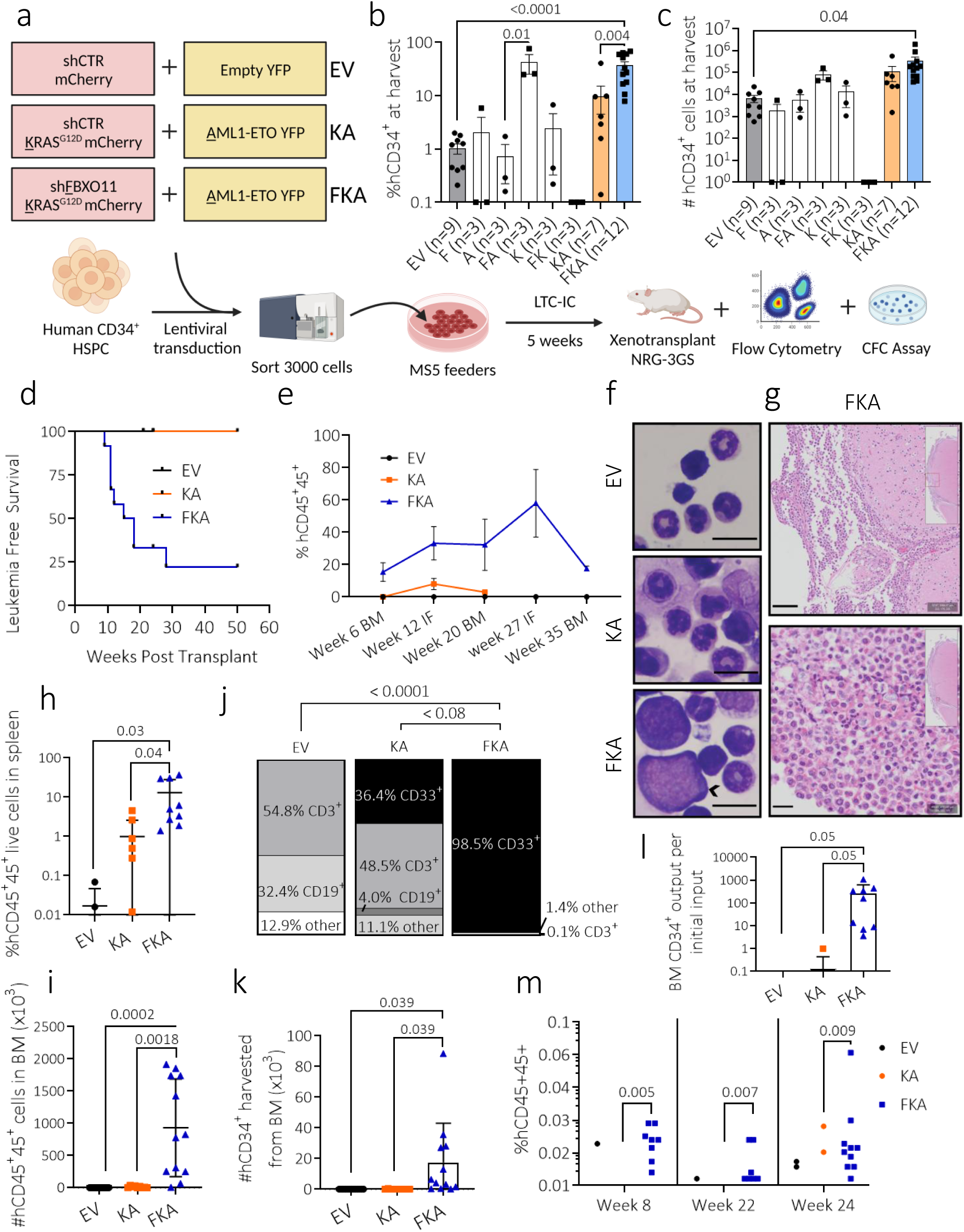
*FBXO11* depletion cooperates with *AML1-ETO* and *KRAS*^G12D^ to generate human myeloid leukemia. **a**, Schematic outlining the experimental design for long-term culture and xenotransplant of CD34^+^ HSPC. (**b**) *FBXO11* knockdown maintains CD34^+^ cells in long-term culture in terms of percentages and (**c**) total live cell counts of CD34^+^ cells were measured on day 40 of culture (*N* = 9 Empty Vector (EV), *N* = 3 sh<u>F</u>BXO11 (F), <u>A</u>ML1-ETO (A), FA, <u>K</u>RAS^G12D^ (K), FK, *N* = 7 (KA), *N* = 12 (FKA) independent replicates). **d**, Kaplan-Meier curve for mice transplanted with cells harvested in (**b**) (*P* = 0.0028 for EV vs FKA, and *P* = 0.0002 for KA vs FKA (Mantel-Cox test)). **e**, Bone marrow engraftment of transduced CD34^+^ cells harvested in (**b**) (EV (*N* = 13); KA (*N* = 9); and FKA (*N* = 12) mice) showing CD45^+^ cell chimerism levels in the non-injected bone (NI) or injected bone (I) (*P* = 0.0033 for EV vs FKA, *P* = 0.018 for KA vs FKA (Sidak’s multiple comparisons test)). **f**, Giemsa-stained bone marrow smear from mice at endpoint shows large immature cells only in FKA marrow indicated with an arrowhead (scale bar = 10 µm). **g,** Representative hematoxylin and eosin stains performed on brain sections of FKA mice with neurologic signs at endpoint indicating myeloid infiltrate (scale bars = 100 µm top, 20 µm bottom). **h**, Human CD45^+^ hematopoietic cell engraftment in the spleen at endpoint as percentage of total viable spleen cells (*N* = 5 (EV, 3 not detected (ND)), *N* = 9 (KA, 1 ND), *N* = 12 (FKA) mice). ND = not detected for values < 0.01%. **i**, Number of human CD45^+^ hematopoietic cell engraftment in the bone marrow measured at endpoint. (*N* = 13 (EV, 7 ND), *N* = 9 (KA), *N* = 12 (FKA) mice). **j**, Percentage of lymphoid (CD3^+^, CD19^+^), myeloid (CD33^+^), and other (CD3^-^ CD19^-^ CD33^-^) cells within the viable human CD45^+^ cell population was measured in bone marrow cells at endpoint. (*N* = 5 (EV), *N* = 5 (KA), *N* = 5 (FKA) mice). **k**, Total CD34^+^ HSPC numbers in the bone marrow were quantified at endpoint. (*N* = 13 (EV, 13 ND), *N* = 9 (KA, 8 ND), *N* = 12 (FKA) mice). **l**, Bone marrow output of CD34^+^ cells per transplanted CD34^+^ cells from LTC-IC (*N* = 13 (EV, 13 ND), *N* = 9 (KA, 8 ND), *N* = 12 (FKA) mice). **j**, Hematopoietic cell engraftment in secondary transplants was assessed at weeks 8, 22 and 24. *P* values represent Fisher’s exact tests. All other *P* values represent two-tailed *t-* tests unless otherwise specified, and error bars represent s.d.

Following long-term culture we xenotransplated EV, KA, and FKA cells into NOD.*Rag1^-/-^;γc^null^* (NRG) mice expressing human *IL3*, human *GMCSF*, and human *SCF* (NRG-3GS) (Figure 7a). FKA, but not KA, xenotransplantion of NRG-3GS mice resulted in significantly reduced overall survival (Figure 7d). However, of the mice that survived, those with transplanted FKA cells had significantly longer and higher human hematopoietic reconstitution compared to KA- and EV-transplanted mice, suggesting that *FBXO11* depletion improves long-term in vivo repopulating ability of human HSPC (Figures 7e and S11a). Morphological examination of FKA mice at endpoint revealed blast cells in the bone marrow, and leptomeningeal infiltration with primitive myeloid cells, consistent with myeloid leukemia (Figures 7f-g and S12). FKA mice had significantly increased human hematopoietic cell reconstitution in the spleen and bone marrow (Figures 7h-i), >98% of which were CD33^+^ myeloid cells similar to human AML (Figures 7j and S13a). At endpoint we also observed a loss of CD3^+^ T cells at the expense of myeloid expansion only in FKA mice (Figures 7j and S13a). FKA-transplanted mice also had greater numbers of CD34^+^ bone marrow cells at endpoint (Figures 7k and S13b). Human CD34^+^ cell output post-transplant per initial CD34^+^ cell placed in long-term culture revealed ∼6,000-fold expansion of FKA HSPC, consistent with the acquisition of self-renewing properties, while KA HSPC did not result in CD34^+^ cell expansion (Figures 7l and Table S6).

Serial marrow transplantation resulted in enhanced reconstitution of human hematopoietic populations with FKA cells compared to controls, indicating maintenance or expansion of self-renewing hematopoietic cells (Figure 6m). Taken together, these results demonstrate that *FBXO11* depletion improves long-term in vivo human cell reconstitution, consistent with induction of a myeloid-skewed, long-term self-renewing stem cell state that cooperates with *KRAS*^G12D^ and *AML1-ETO* to generate myeloid leukemia.

## DISCUSSION

In this work we highlight ubiquitin pathway mutations as a novel class of AML mutations. We implicate *FBXO11* depletion in leukemogenesis, and identify a novel connection between ubiquitination and mitochondrial function through FBXO11-mediated regulation of the mitochondrial protease and chaperone, LONP1, independent of the proteasome system.

Dysregulation of LONP1 has been associated with colon cancer and melanoma, but has not been studied in the context of AML^33,41,42^. Consistent with previously characterized activating effects of K63-linked polyubiquitination of target proteins, our data show that FBXO11 ubiquitinates LONP1 to relocate LONP1 into the mitochondria and regulate respiration. This activating effect of FBXO11 contrasts with other FBXO11 targets previously reported. For instance, SCF^FBXO11^ has been shown to target BCL6, CRL4, and Snail family transcription factors for ubiquitination and subsequent proteasomal degradation^43–45^. SCF^FBXO11^-mediated neddylation of p53 also inhibits p53 transcriptional activity^35^. Our findings provide an explanation as to why *LONP1* was not detected in a screen for identifying mitochondrial contributors to leukemogenesis, as the readout in that study was reduced leukemia cell viability with target gene knockdown^46^. In our model, it is reduced LONP1 function due to *FBXO11* depletion, not LONP1 activation, that is responsible for leukemogenesis.

Transcriptomic and functional data support FBXO11 as an upstream regulator of LONP1 and solidify the link between *FBXO11* and *LONP1* depletion in regulating mitochondrial function. The mitochondrial respiratory chain defects are consistent with the reduced MMP observed in long-term self-renewing stem cells^23–25,47,48^. Of interest, the reduction in erythroid differentiation resulting from *LONP1* depletion, which mimicked the effects of *FBXO11* depletion and is also reflected in the transcriptomic data, is supported by a previous report on the role of LONP1 in heme biosynthesis^49^.

Loss of ETC function can result from the accumulation of misfolded ETC proteins shown to occur with *LONP1* loss, or from loss of LONP1 chaperone activity required for proper folding of some ETC proteins^32,50^. We found that both *FBXO11* and *LONP1* depletion reduced MMP, and that FBXO11 acts through by K63-linked ubiquitination and regulation relocalization of LONP1 into the mitochondria. LT-HSCs and CD34^+^ cells that have greater reconstitution potential are both characterized by low MMP^24,26,34^. Disruption of MMP also promotes self-renewal in HSCs, indicating that low MMP is essential for maintaining an HSC state^24^. While *FBXO11* depletion has previously been shown to impart apoptotic effects^51^, our work shows that this is not true for primitive HSPC. Rather, *FBXO11*-depleted HSPC are protected from apoptosis and remain in G_0_ of the cell cycle. These findings are also consistent with the stem cell characteristics we observed with *FBXO11* depletion, both functionally, and at the transcriptional level. Indeed, the stem cell phenotype induced by *FBXO11* depletion is sufficient to cooperate with *AML1-ETO* and *KRAS*^G12D^ to generate a synthetic serially-transplantable AML in normal CD34^+^ cells.

While mitochondrial metabolism and function in stem cells have recently been a focus in stem cell research, there is no consensus on how exactly reduced mitochondrial function promotes a stem cell state or contributes to leukemogenesis. Metabolic changes and products of metabolic pathways have been shown to impact nuclear transcription as well as chromatin and epigenetic remodeling, particularly in cancers^52,53^. In fact, changes in MMP have previously been shown to affect nuclear gene expression in macrophages, suggesting one mechanism for how reduced mitochondrial function might promote a stem cell state^54^.

In this work, we suggest that mutations of ubiquitin pathway genes may represent a novel molecular subclass in AML. We draw a novel connection between the ubiquitin pathway and mitochondrial respiration. In particular, we show that LONP1 is a target that is activated, rather than degraded following FBXO11-mediated ubiquitination, and this acts as a regulator of hematopoiesis. Importantly, we show that depletion of *FBXO11* cooperates with *AML1-ETO* and activated *KRAS*^G12D^ to generate serially transplantable leukemia in a xenograft model. This work suggests that altered gene expression independent of genetic variants may implicate ubiquitin system gene defects in a much wider cohort of AML patients.

### Limitations of the study

While a number of ubiquitin system gene mutations were identified we only characterized the role of a single recurrently mutated gene in AML. Hence, the impact of other ubiquitin gene mutations on AML pathogenesis would need to be examined independently. Although ubiquitin system gene mutations were found to be mutually exclusive, it remains to be understood whether this creates a common dependency or vulnerability for ubiquitin system genes, or whether there is synthetic lethality when multiple ubiquitin genes are mutated in the same cell. While the data support LONP1 as a primary ubiquitination target influencing the phenotype observed in AML, these studies do not exclude the influence that other FBXO11 targets may have in AML pathogenesis. Loss of FBXO11 prevents relocalization of LONP1 into the mitochondria, but we were not able to enforce mitochondrial relocation of LONP1 in the absence of FBXO11. Hence the relationship of FBXO11 on LONP1 localization and function could only be examined in the context of FBXO11 overexpression.

## Acknowledgements

The authors would like to thank all members of the Karsan laboratory for their comments and feedback. We would like to acknowledge the patients and their families who provided samples. This work was supported by grants to AK from the Canadian Institutes of Health Research (PJT-162131 and PJT-183924) and the Terry Fox Research Institute (Project #1074). H.K. is supported by a Killam Doctoral New Scholar Award. A.K. is a Tier 1 Canada Research Chair in Myeloid Cancers. The authors would also like to thank Maria Koh for assistance with designing the graphical abstract.

## Author Contributions

A.Y.C.M., H.K., X.W., L.Y.T.C., G.D., F.M., Z.Z., J.T., N.G., J.J., J.P., and R.K.G. contributed to the performance of the experiments and/or analysis of the data. H.K., T.L., T.R.D., S.H., E.W. and J.B curated data and performed bioinformatics assays. S.C. and S.W.G.C. performed mass spectrometry analyses. D.W. performed and interpreted pathology on murine specimens. R.K.G., G.B.M., and A.K. provided advice and interpreted the data. A.K. conceived, secured funding, and supervised the project. A.Y.C.M., H.K., and A.K. wrote the manuscript.

## Declaration of interests

The authors declare no competing interests.

## METHODS AND MATERIALS

### Mutation calling for oncoprint inclusion

Patient mutations were determined pathogenic and included in the oncoprint if they had a REVEL score > 0.250 and CADD score > 10.5. Germline variants were excluded. Hugo Gene Nomenclature Committee (HGNC) gene data was used to generate a list of ubiquitination genes to query, and the list was manually filtered to retain genes functionally relevant to the UPS. For determining the REVEL score threshold of 0.25 for pathogenicity, data for pathogenic solid tumor variants were obtained from *Cheng et al* (Memorial Sloan Kettering-Integrated Mutation Profiling of Actionable Cancer Targets (MSK-IMPACT))^55^, and variants that were reported less than five times in the Catalogue of Somatic Mutations in Cancer (COSMIC) were excluded. Pathogenic variants associated with human hematological cancers were obtained from *Jaiswal et al* ^56^, and variants that were reported less than 5 times in COSMIC were excluded.

### Odds ratios calculations

Odds ratios were calculated as “odds of a mutation occurring in a sample with an ubiquitin pathway mutation” vs “odds of a mutation occurring in a sample without ubiquitin pathway mutations”. Odd ratio for mutations in gene

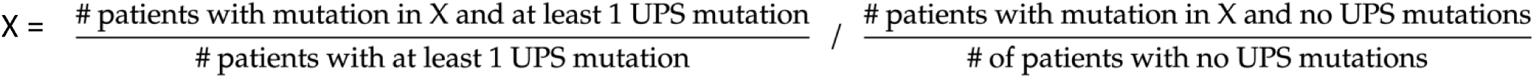

### Isolation of human HSPCs

Human HSPC isolation was performed by the Stem Cell assay core at BC Cancer Research Institute. Anonymized normal human cord blood samples were obtained with informed consent from normal full-term Cesarean section deliveries in accordance with procedures approved by the Research Ethics Board of the University of British Columbia and samples from a single day pooled for further processing. CD34^+^ HSPCs were obtained at >95% purity from pooled collections using a two-step Rosette-Sep/EasySep human CD34-positive selection kit (STEMCELL Technologies) according to the manufacturer’s protocols and/or FACS sorting. CD34^+^ HSPC cells were thawed by dropwise addition into HBSS + 10% fetal bovine serum (FBS) supplemented with DNase (100 μg/mL final concentration). CD34^+^ HSPCs were then seeded into 96-well round bottom plates (cell density at 100,000 (100k) cells/100 μL) and prestimulated in StemSpan^TM^ SFEM (Stem Cell Technologies #09650) supplemented with StemSpan™ CC100 (Stem Cell Technologies #02690), 750 nM SR1 (Stem Cell Technologies #72344) and 35 nM UM171(Stem Cell Technologies #72912) overnight.

### Cell lines

HEK293T, K562, HL60, and AML-193 were purchased from the American Type Culture Collection. MOLM-13, Kasumi-1, OCI-AML3, OCI-AML2, and OCI-AML5 were purchased from the Leibniz Institute DSMZ. HEK293T were maintained in Dulbecco’s modified Eagle’s medium (DMEM) supplemented with 10% fetal bovine serum (FBS). OCI-AML2 was cultured in α Minimum Essential Medium supplemented with 10% FBS. K562, HL60, MOLM-13, OCI-AML3, OCI-AML5, and AML-193 cell lines were cultured in RPMI 1640 medium supplemented with 10% FBS, or 20% FBS for Kasumi-1. Media were supplemented with 2 ng/ml GM-CSF (BioLegend 572902) for OCI-AML5, or 0.005 mg/ml insulin (Sigma-Aldrich 19278), 0.005 mg/ml transferrin (Sigma-Aldrich T1147), and 5 ng/ml GM-CSF for AML-193. All of the cell lines were confirmed by single-nucleotide-polymorphism array and short-tandem-repeat authentication (Genetica DNA Laboratories) and routinely tested negative for mycoplasma (MycoAlert, Lonza)

### shRNA constructs and CRISPR clones

K562 control and *FBXO11* knockout CRISPR clones were generated by Dr. Gerben Duns. Guide RNAs (gRNAs) were cloned into the pSpCas9n(BB)-2A-GFP (PX461) plasmid (Addgene #48140) and delivered to parental K562 cells by Amaxa nucleofection using the Amaxa Cell line nucleofector kit V (Lonza VCA-1003). GFP^+^ transfected cells were single-cell sorted into 96-well plates, expanded. *FBXO11* targeting was confirmed by Sanger sequencing, and FBXO11 depletion was confirmed by western blot. shRNAs used are in the pLKO.1 vector, from the MISSION library, and received from the UBC Centre for High-Throughput Biology, and GFP or mCherry was inserted in place of the puromycin resistance marker. gRNA and shRNA information are listed below. More information for gRNA and shRNA targeting can be found in Appendix Figures A.3 and A.4.

*<u>FBXO11</u>* <u>CRISPR gRNAs</u>

gRNA sequence: AACCACTGTAGGGTTAGCATAGG
Target exon: Exon 12/23 (antisense, positive strand)

<u>Human shRNAs</u>

shCTR
Target sequence: CCTAAGGTTAAGTCGCCCTCGC

sh*FBXO11* #9
TRC Clone ID: TRCN0000004303
Target exon: Exon16/23 (isoform 4 NM_001190274.2)
Target sequence: GAGTGCTAGAAGACAATGATA

shFBXO11 #10
TRC Clone ID: TRCN0000004304
Target exon: Exon 5/23 (of isoform 4 NM_001190274.2)
Target sequence: GAGAGTTTCCAGCAGTTGTAT

shLONP1 #1
TRCN0000046793, Exon 11/18 (isoform 1 NM_004793.4)
Target sequence: CCAGTGTTTGAAGAAGACCAA

shLONP1 #3
TRCN0000046797 Exon 2/18 (isoform 1 NM_004793.4)
Target sequence: CCAGCCTTATGTCGGCGTCTT

### Lentiviral constructs and transduction

High-titre lentiviral supernatants were produced by transient transfection of 293T/17 cells with second-generation packaging/envelope vectors pRRE (Addgene #12251), pREV (Addgene #115989), and pMD2.g (Addgene #12259) using CalPhos^TM^ mammalian transfection kit (Clontech #631312) and incubated for 16 hours, followed by ultracentrifuge concentration (25,000 rpm for 90 min at 4°C; Beckman SW32Ti rotor).

CD34^+^ HSPCs were transduced in 96-well plates (Thermo Scientific #12-565-65) with concentrated lentivirus. After 6 hours of incubation, cells were washed once with PBS and transferred to culture medium (StemSpan^TM^ SFEM with StemSpan™ CC100 to culture medium at a 1 in 100 dilution, 750 nM SR1 and 35nM UM171). Following 72-hour ex vivo culture, transduced cells were then isolated by sorting for reporter fluorophore expression from lentiviral constructs on a Fusion sorter (Becton Dickenson) using purity mode.

### *Ex vivo* cord blood liquid culture for differentiation assay

Transduced cells were sorted into 96-well plates at 10k cells/well into 100 μL in StemSpan^TM^ SFEM supplemented with StemSpan™ CC100, 20 ng/mL recombinant human thrombopoietin (hTPO) (BioLegend #597402), DNase (100 μg/mL final concentration), and antibiotics (penicillin and streptomycin). Cells were incubated in a humidified atmosphere at 37°C in room air supplemented with CO2 to achieve a final concentration of 5%.

### Cell differentiation analysis

CD34^+^ HSPCs were harvested from culture and washed with PBS with 2% FBS before staining with antibodies at the dilutions listed in the Table below for 1 hour on ice. Cells were pelleted at 300 x g for 5 min, resuspended in PBS with 2% FBS and 1 μg/mL DAPI, and filtered before flow cytometry analysis.

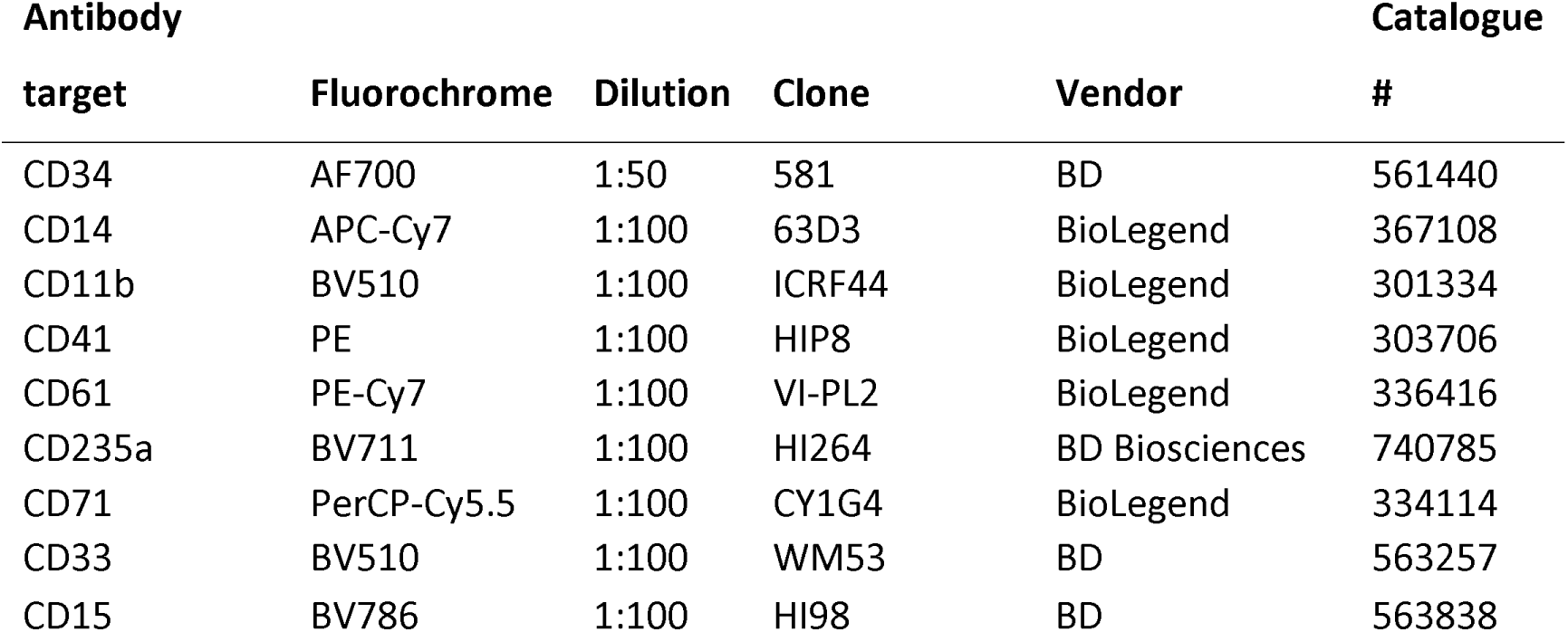

### Long-term initiating cell (LTC-IC)

Two-three days before cell sorting for long-term co-culture with MS5 feeder cells, frozen and X-ray irradiated feeder cells were thawed via dropwise addition into 9 mL DMEM with 15% FBS (15% DMEM) (Invitrogen 12100-046). The cell mixture was spun down at 300 × g for 5 min and the supernatant was decanted. The cell pellet was then resuspended in 6 mL 15% DMEM (no antibiotics) and plated at 1 mL/dish in collagen-coated 35 mm dishes (collagen was neutralized with DMEM medium before cell plating). The feeder cells were incubated in a 37° C incubator overnight. The media was replaced with 2 mL fresh 15% DMEM medium the next day. On the day of CD34^+^ HSPC cell sort, the media was removed and replaced with 2.5 mL of LTC media (STEMCELL Technologies #05150) supplemented with 10 μM Hydrocortisone (STEMCELL Technologies #07904).

Weekly half-medium changes were performed by removing one half of the medium containing cells and replacing with freshly prepared H5100 with 10 μM hydrocortisone over the 40-day culture period.

### Colony-forming cell (CFC) assay

For primary CFCs, 500 of sorted lentivirally transduced CD34^+^ HSPCs were seeded into 1 mL of MethoCult^TM^ H4434 Classic media (STEMCELL Technologies #04434) supplemented with StemSpan™ CC100 with antibiotics, and plated in 35mm CFC dishes (STEMCELL Technologies #27150). Cells were incubated for 10 days before colonies were counted.

For secondary CFCs, the media containing the cells were collected from the CFC dishes and transferred into 15mL tubes. PBS with 2% FBS was used to rinse the plates and pooled with the collected media. Cells were spun down at 300 x g for 5min, washed with PBS with 2% FBS. After cell counting, the cells were seeded into fresh CFC dishes in fresh MethoCult^TM^ H4434 Classic media supplemented with StemSpan™ CC100 and antibiotics at 50k cells/mL in each plate. Colonies were counted after 10 days.

### Xenotransplantation

All experimental procedures were done in accordance with protocols approved by the University of British Columbia Animal Care Committee. NRG-3GS mice (NOD.Cg-*Rag1tm1Mom Il2rgtm1Wjl* Tg(CMV-IL3,CSF2,KITLG)1Eav/J)) were bred and maintained in the Animal Resource Centre of the British Columbia Cancer Research Centre under pathogen-free conditions.

Female NRG-3GS mice 8-15 weeks of age were sublethally irradiated with 850 cGy (cesium irradiation, medium dose rate) on the day of intrafemoral injection. 100,000 cells harvested from LTC were injected per mouse. For transplant analysis of *ex vivo* cultured cells, bone marrow aspirates were performed every 6 weeks post-transplant and human chimerism was tracked by flow cytometry analysis (using a BD LSR Fortessa analyzer or BD FACSymphony) and analyzed with the following: APC-anti-CD45 2D1(1:50; Invitrogen 17-9459-42), APC/Cy7 anti-CD45 HI30 (1:50; BioLegend 304014), AF700-anti-CD34 (1:50; BD biosciences 561440), PE/Cy7-anti-CD33 (1:100; Thermo fisher 25-0338-42), PE/Cy7-anti-CD15 (1:100; BD biosciences 560827), BV650-anti-CD19 (1:100; BD biosciences 560827), SB600-anti-CD3(1:50; Invitrogen 83-0037-42), DAPI (1 μg/mL final concentration, Sigma-Aldrich D9542).

Bone marrow aspirates were performed on the non-injected femur (BM, weeks 6, 20, and 35 weeks post-transplant) or the injected femur (IF, weeks 12 and 27 weeks post-transplant) to determine human cell chimerism. Engraftment is considered long term if detection threshold of >0.01% CD45^+^ cells is exceeded at 20 weeks.

Transplanted mice were sacrificed at predefined, humane clinical morbidity endpoints. At endpoint, human chimerism in bone marrow (cells flushed from femurs, pelvis and tibias), spleen, and liver were tracked by flow cytometry analysis (using a BD LSR Fortessa analyzer or BD FACSymphony) and analyzed with the following: APC-anti-CD45 2D1 (1:50; Invitrogen 17-9459-42), APC/Cy7-anti-CD45 HI30 (1:50; BioLegend 304014), AF700-anti-CD34 (1:50; BD biosciences 561440), PE-anti-CD33 (1:50; eBioscience 8012-0337-120), PE/Cy7-anti-CD15 (1:100; BD biosciences 560827), BV480-anti-CD14 (1:50; BD biosciences 566190), DAPI (1 μg/mL final concentration, Sigma-Aldrich D9542).

For secondary transplants, viably frozen bone marrow harvested at primary transplant endpoints were thawed and added drop-wise into 8mL thawing medium (PBS with 10% FBS and 100 μg/mL DNAse final concentration). Cells were spun down at 400 x g for 5 min to remove the supernatant. Cells were counted, and the total amount of harvested bone marrow for each EV and KA mice were transplanted into each sublethally irradiated recipient NRG-3GS by tail vein injection. Half of the harvested bone marrow from each donor FKA mouse was transplanted into each FKA recipient mouse. Secondary transplanted mice were monitored, processed, and analyzed following the procedures for primary transplants.

### Calculating CD34^+^ cell output for xenotransplants

The number of human CD34^+^ cells generated per input CD34^+^ cell after the LTC-IC and xenotransplant process was calculated based on the assumption below that: the ratio of the number of CD34^+^ cells harvested from the LTC (**A**) to the number of CD34^+^ cells that were injected into the mice (**B**) equaled the number of CD34^+^ cells that would have been harvested at xenotransplant endpoint if the total amount of CD34^+^ cells harvested from the LTC were injected into the mice(**C**) to the number of CD34^+^ cells that were actually harvested at xenotransplant endpoint (**D**). Since equal numbers (100,000) of cells harvested from LTC were injected into recipient mice for xenotransplantation, the number of CD34^+^ transplanted varied between conditions (Supplementary Table 6).

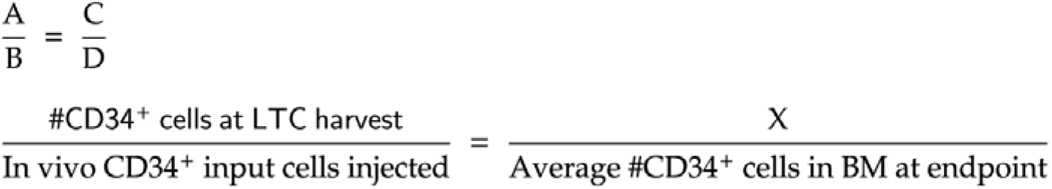

To calculate the number of CD34^+^ cells generated by the end of the experiment per CD34^+^ cell input into the LTC-IC experiment, we divided “X” by the number of LTC-IC input CD34^+^ cells, which was the same for each condition.

### RNAseq extraction and RNAseq

CD34^+^ HSPCs were cultured for 24 hours in StemSpan^TM^ SFEM supplemented with StemSpan™ CC100, 20 ng/mL recombinant hTPO, 750 nM SR1, 35 nM UM171, DNase (100 ug/mL final concentration), and antibiotics after sorting for lentivirally transduced cells as described above before pelleting at 300 x g for 5 min. Cell pellets were resuspended in TRIzol^TM^ reagent (ThermoFisher #15596026). Total RNA was extracted according to manufacturer guidelines. For RNA precipitation, 5 μg glycogen (Roche #10901393001) was added per sample. 40 ng of RNQA per sample was submitted based on quantification by NanoDrop^TM^. Downstream processing, quality control, and RNAseq was performed at Michael Smith laboratories facilities by 1 x Illumina seq with 150bp PET Indexed Lanes. cDNA was labelled using the Biotin Allonamide Triphosphate protocol. Target hybridization was performed at 45°C, for 16-18hr using the FS450_0001 fluidics protocol for washing. Data was acquired using the Affymetrix GeneChip Scanner 3000. CEL files were imported into R and the probe expression values were normalized using the Robust Multichip Average (RMA) method from the ’oligo’ (v.1.50.0) package. Probes that were lowly expressed, defined as not having normalized expression values greater than 4 in at least 3 samples, were filtered out. Probes that ambiguously mapped to multiple genes were filtered out.

### Ki-67 staining

CD34^+^ HSPC were fixed and permeabilized with the BD Fixation and Permeabilization kit (#554714) according to manufacturer protocol. Fixed and permeabilized cells were stained with PE anti-human Ki-67 antibody (BioLegend #350504) diluted 1:10 in the BD Perm/Wash buffer from the kit for 30 min on ice. Unbound antibody was washed with the Perm/Wash buffer. Pelleted cells were resuspended in BD Perm/Wash buffer with 2 μg/ml DAPI and incubated for 10 min in the dark, on ice. Cells were centrifuged at 1500 x g for 5 min and resuspended in PBS with 2% FBS and strained for flow cytometry analysis.

### Histology

Mouse brain samples were fixed in 10% buffered formalin and stored at 4°C for a week before paraffin embedding. H&E and myeloperoxidase staining of brain samples were performed at the BC Cancer Vancouver Center Pathology department. Bone marrow smears were stained with Wright-Giemsa staining solution.

### Immunoprecipitation

For co-immunoprecipitations for identifying protein-protein interactions, cells were washed in PBS then lysed with modified RIPA buffer (50 mM Tris-HCl pH 7.6, 150 mM NaCl, 1 mM EDTA, 1% Triton X-100, protease inhibitor cocktail 1 tablet/1 mL at 1:100 (Sigma-Aldrich 11697498001), phosphatase inhibitor cocktail 1:100 (Sigma-Aldrich P0044-5ML), 1 mM Na3VO4 and 10 mM NaF) for 1 hour at 4°C on a rotating platform, then passed 10x through a 22-G needle. The lysate was centrifuged at 16,000 x g for 10 min at 4°C. The supernatant was collected and protein quantification was performed using the BCA assay (BioRad DC Protein assay kit, Bio-Rad 5000112). 1 mg of protein per condition was used for co-immunoprecipitation, and the volume was made up to 1 mL with the lysis buffer. The lysate was precleared with Protein G agarose beads (EMD Millipore 16-266) for FLAG immunoprecipitation or Protein A agarose beads (EMD Millipore 16-125) for LONP1 immunoprecipitation on a rotating platform at 4°C, overnight (16 hours) for FLAG immunoprecipitation or for 3 hours for LONP1 immunoprecipitation. Beads were spun down at 1500 x g for 1 min, and the lysate was transferred to a fresh tube with 50 μL equilibrated ANTI-FLAG® M2 Affinity Gel (Sigma-Aldrich A2220-1ML) for FLAG immunoprecipitation, or with Protein A agarose beads incubated with 2 μg anti-LONP1 antibody (ThermoFisher Scientific 15440-1-AP) overnight (16 hours) and incubated on a rotating platform at 4°C for 1 hour for FLAG immunoprecipitation, and for 3 hours for LONP1 immunoprecipitation. The beads were washed with wash buffer (50 mM Tris-HCl pH 7.6, 150 mM NaCl, 0.05% Triton X-100) 5 times with 1 min 1500 x g spins in between, and eluted in 1x Laemmli sample buffer at 95°C for 15 min.

For LONP1 immunoprecipitation to identify post-translational modifications, cells were lysed with RIPA buffer (25mM Tris, 150mM NaCl, 1% NP40, 0.5% sodium deoxycholate, 0.1% sodium dodecyl sulfate (SDS) instead of modified RIPA buffer.

### Quantitative tandem mass spectrometry

K562 cells lentivirally expressing FLAG-*FBXO11* were lysed in modified RIPA buffer with 10 μM MG132 (ApexBio A2585-25) as described above. Whole cell lysate was incubated with Anti-FLAG® M2 Magnetic Beads (Sigma-Aldrich M8823-1ML) or IgG beads for 3 hours then washed with lysis buffer 3 times. Three replicates per condition were directly eluted with SP3 elution buffer (200 mM HEPES pH 8.0, 10% SDS, 200 mM DTT). Four replicates each were treated with Benzonase® (EMD Millipore 70664-3) before elution. TMT labelling was performed by the mass spectrometry core at the Genome Sciences Centre. 8-plex TMT labels were used. Quantification results were calculated based on published methods^57,58^.

For the set of immunoprecipitations performed without Benzonase®, proteins had normalized protein enrichment score (PESn) greater than 0.6, and were enriched in 2 of the 3 replicates by at least 1.1-fold were shortlisted. For the immunoprecipitations performed with Benzonase® treatment, proteins that had PESn greater than 0.45 and were enriched in 3 of 4 replicates by at least 1.2-fold were shortlisted. The shortlisted proteins from each experiment were intersected to identify commonly immunoprecipitated proteins.

### Measuring mitochondrial membrane potential

Mitochondrial membrane potential (MMP) was measured by TMRE for cell lines, primary AML samples and CD34^+^ HSPC (Cedarlane labs ENZ-52309), or with the JC-1 dye (eBioscience 65-0851-38) for CD34^+^ HSPC (Figure 5f). For TMRE, cells were cultured in their regular culture media and conditions with 400 nM TMRE for 30 min. Cells were spun down at 400 x g for 5 min to remove the media and TMRE, and the cell pellets were resuspended in PBS with 2% FBS and 1 μg/mL DAPI and kept on ice before analysis by flow cytometry. JC-1 dye was used according to manufacturer protocol. Stained cells were resuspended in PBS with 2% FBS and 1 μg/mL DAPI for flow cytometry analysis.

### Primary AML samples and CD34^+^ HSPC media for measuring mitochondrial membrane potential

Primary AML samples and CD34^+^ HSPC were cultured in medium previously described^59^. STEMdiff APEL2 (STEMCELL Technologies 05275), 2% FBS (Sigma-Aldrich F1051), 2% Chemically Defined Lipids (Thermo Scientific 11905031) supplemented with 25 ng/mL VEGFA (STEMCELL Technologies 78159.1), 25 ng/mL VEGFC (STEMCELL Technologies 78202.1), 25 ng/mL bFGF (STEMCELL Technologies 78134.1), 25 ng/mL hSCF (Stem Cell Technologies 78062), 25 ng/mL Flt3 (Stem Cell Technologies 78009) and 10 ng/mL TPO (STEMCELL Technologies 78210.1), 10 ng/mL EPO (STEMCELL Technologies 78007), 10 ng/mL IL3 (STEMCELL Technologies 78040.1), 10 ng/mL IL6 (Cedarlane Labs 206-IL-200/CF). 50% media change was performed on day 3, 24h before TMRE staining.

### Mitochondrial mass measurements

MitoTracker™ Deep Red FM (Fisher Scientific M22426) was used to measure mitochondrial mass in HSPC. Cell staining was performed according to the manufacturer’s protocol. Cells were stained for 30 min, then washed with PBS with 2% FBS and resuspended in PBS with 2% FBS and 1 μg/mL DAPI for flow cytometry analysis.

### Ubiquitin, neddylation activating enzyme, and proteasome inhibitor treatment

Ubiquitin activating enzyme (UAE) inhibitors PYR-41 (Sigma-Aldrich N2915-5MG) was used at 50 nM, and TAK-243 (also known as MLN7243, Selleck Chemicals S8341) at 0.1 μM. Neddylation activating enzyme (NAE) inhibitors MLN4924 (Cayman Chemical Company 15217) was used at 50 nM, and TAS4464 (Selleck Chemicals 15217) was used at 50nM. Proteasome inhibitors MG132 (ApexBio A2585) was used at 1μM, and Bortezomib (Selleck Chemicals S1013) at 50nM. Concentrations were determined by published concentrations used on K562 cells or other myeloid cell lines, and inhibitory effect on global ubiquitination or neddylation levels were confirmed by western blot. Cells were seeded at 300k cells/mL and treated with the above inhibitors added directly to culture media for 16 hours. Inhibitor stock solutions were prepared in DMSO.

### Western blots

Cells were washed with PBS and lysed for 30 min on a rotating platform at 4°C in RIPA buffer (25 mM Tris, 150 mM NaCl, 1% NP40, 0.5% sodium deoxycholate, 0.1% sodium dodecyl sulfate (SDS), protease inhibitor cocktail, phosphatase inhibitor cocktail, 1 mM Na3VO4 and 10 mM NaF). Lysed cells were centrifuged for 16,000 x g for 10 min at 4°C, and the supernatant was collected for protein quantification by BCA. Samples were mixed with Laemmli sample buffer incubated at 95°C on a heat block for 15 min.

Proteins were separated by SDS-polyacrylamide gel electrophoresis at 120 V at room temperature and transferred to nitrocellulose membrane at 100 V at room temperature for 1 hour for proteins of interest under 55 kDa, and at 35 V at 4°C for 16 hours for proteins of interest over 55 kDa. After transfer, blots were rinsed briefly with tris-buffered saline with 0.1% Tween 20 (TBST), and then blocked for 1 hour at room temperature in 5% skim milk in TBST. Membranes were washed 3 times for 10 minutes each with TBST.

Blots were probed with primary antibodies (FBXO11 antibody at 1:1000 (Bethyl Laboratories A301-178A), GAPDH antibody at 1:10,000 (Sigma-Aldrich G8795-200UL), HSP60 1:1000 (Cell Signaling Technology 4870S), LONP1 antibody at 1:1000 (ThermoFisher Scientific 15440-1-AP), K63 linkage specific antibody 1:1000 (Cell Signaling Technology 5621S), NEDD8 antibody at 1:1000 (Cell Signaling Technology 2745S), α-tubulin at 1:1000 (Sigma-Aldrich T5168-.2ML) TOM20 at 1:1000 (New England Biolabs 42406S) were prepared in TBST with 3% bovine serum albumin (BSA), based on manufacturer recommended incubation times. Membranes were washed 3 times for 10 minutes each with TBST and probed with appropriate HRP secondary antibody (anti-Rabbit Immunoglobulin/HRP, Dako P044801 or anti-mouse Immunoglobulin/HRP, Dako P0260) at 1:1000 in TBST with 5% skim milk. Blots were developed with Western Lightning Plus, Chemiluminescent Substrate (PerkinElmer NEL105) and imaged with a BioRad ChemiDoc MP Imaging system.

### Mitochondria Fractionation and isolation by differential centrifugation

Mitochondria were isolated from *FBXO11*-KO and OCI-AML3 cells as described^60^, with the protocol scaled down for 1 x 10^8^ cells. Briefly, cells were pelleted and washed with PBS before being swollen for 10 minutes with hypotonic buffer and ruptured by using a dounce homogenizer. The homogenized supernatant was centrifuged 3 times at 1300 x g for 5 minutes, and then at 17,000 x g for 15 minutes to pellet the mitochondria. Mitochondria were washed once and pelleted again, before being lysed with RIPA buffer as described above.

### Immunofluorescence staining, proximity ligation assay, and confocal microscopy

Coverslips (Fisher Scientific 12-545-81) were coated with 0.01% poly-L-lysine solution (Sigma-Aldrich P8920-100ml) for 2 hours, washed twice with ddH_2_O, and dried overnight. Cells were incubated in fresh medium for 1 hour on the poly-L-lysine coated coverslips before being washed with PBS, fixed with 3% paraformaldehyde, and permeabilized with 0.01% Triton-X. Samples were blocked with PBS supplemented with 1% BSA, 10 mM MgCl_2_, and 1 mM CaCl_2_ for 1 hour at room temperature. Samples were stained with primary antibodies to FBXO11 (1:50, Abnova H00080204-M01) and LONP1 (1:100,

Thermo Fisher 15440-1-AP) for 2 hours, and secondary antibodies (1:500 goat anti-mouse Alexa Fluor 488, Fisher Scientific A11008 / goat anti-rabbit Alexa Fluor 546, Fisher Scientific A11035) for 1 hour at room temperature.

The Duolink Proximity Ligation Assay (PLA) was performed according to the manufacturer’s protocol. Instead of secondary antibody staining described above, samples stained with primary FBXO11 (mouse) and LONP1 (rabbit) antibodies were incubated with Anti-Rabbit PLUS (Sigma-Aldrich DUO92002) and Anti-Mouse MINUS (Sigma-Aldrich DUO92004) *in sit u*probes, ligated and amplified using red detection reagents (Sigma-Aldrich, DUO92008).

After final washes following Alexa fluor-conjugated antibody staining or PLA, all samples were co-immunostained with Alexa Fluor 647 conjugated TOM20 antibody (1:1000, Abcam ab209606) for 2 hours at room temperature to identify mitochondria. All samples were then mounted on slides with ProLong Gold Antifade Mountant with DAPI (Thermo Fisher P36935).

All samples were imaged on a Leica SP5 laser scanning confocal on 63X/1.4 oil objective and processed in ImageJ (Fiji). Colocalization was determined by measuring the area of fluorescence for the indicated proteins and their overlap area in single cells using composite RGB images.

### Seahorse XF Cell MitoStress Test

*FBXO11-*KO clone cells were in the growth phase at the time of the assay. The cells were counted and seeded at 80,000 cells per well in the 96-well Agilent Seahorse XF Cell Mito Stress Test (Agilent 103010-100) assay plate. The assay was performed as outlined in the manufacturer’s protocol and run on the Seahorse XFe bioanalyzer. Analysis and interpretation were done according to the manufacturer’s protocol.

### BloodSpot data analysis

Transcriptomic data for SCF genes from the BloodSpot database (https://servers.binf.ku.dk/bloodspot/) were taken from GSE42519 and GSE13159 datasets. Data for *FBXO11* expression taken from the same datasets, with data for the 219208_at probe.

### RNAseq analysis

RNAseq read alignment was performed using STAR aligner, and count generation was done with Salmon. Variance stabilizing transformation was performed to allow comparison between samples.

Transcript per million (TPM) values were filtered to remove genes that have low counts (TPM < 0.5, equivalent to a count of 10 for the dataset) for at least 2 samples. Differential gene expression analysis was performed on the RNAseq data using DeSeq2 package with R. Gene Set Enrichment Analysis was performed using the Hallmark, C2, and C5 gene set collections from MSigDB. GSEA was performed using pre-ranked gene lists for each comparison presented, where the gene list was ranked based on the log2 fold change value. CellRadar plots from Figure 6 were generated using the public user interface provided by the Karlsson lab (https://karlssong.github.io/cellradar/). The same definition of differentially expressed genes was used, and HSPC data was taken from the BloodSpot HemaExplorer human normal hematopoiesis dataset.

### Deconvolution of AML sample hierarchies from bulk RNAseq and stratification by *FBXO11* expression

Deconvolution of 864 AML sample hierarchies were performed using the original samples and code provided by Zeng et al^38^ (https://github.com/andygxzeng/AMLHierarchies). Following deconvolution and projection by Principal Component Analysis (PCA) patients were stratified into tertiles within their datasets (TCGA, BEAT-AML, Leucegene) based upon their *FBXO11* expression, and their tertile identifier were projected onto the PCA. The sample’s original hierarchy from Zeng et al. was identified for each *FBXO11* expression tertile.

### Statistical Analysis

Measurements for experiments were taken from distinct samples. All statistical tests were performed using GraphPad Prism version 8.0 (San Diego, California). All data are presented as mean ± standard error of the mean (SEM). Statistical significance was set at a threshold of P < 0.05 and FDR < 0.25.

## Supplemental Information

Document S1: Figures S1-S13.

**Supplementary Figure 1. Mutations affecting the ubiquitin pathway occur frequently in AML.**

**a**, REVEL scores for different mutation datasets were compared to identify a threshold for pathogenicity prediction. Mutations plotted include: (1) Known actionable pathogenic missense variants in solid tumors (*n* = 146 variants). (2) Pathogenic variants associated with human hematological cancers (Jaiswal)^56^ (*n* = 149 variants). (3) Somatic missense mutations in AML identified in TCGA^21^ (*n* = 1233 mutations). Applying a minimal REVEL threshold of 0.250, represented by the red dotted horizontal line, removes more than half of all identified somatic missense TCGA variants, while retaining 98% of known pathogenic variants. **b**, Single nucleotide, indel and copy number variants (CNV) affecting the SCF*^FBXO^*^11^ complex are summarized in the oncoprint. In the combined AML PMP, TCGA, Beat AML and TARGET pediatric AML datasets, 7.1% of all 1727 AML patients (123/1727) had a mutation or copy loss of a gene affecting the SCF*^FBXO11^* complex.

**Supplementary Figure 2. *FBXO11* is recurrently mutated in multiple cancers.**

**a**, *FBXO11* mutation types, locations, and frequencies identified in all TCGA cancer samples (n = 10,753) excluding AML samples are shown (cBioPortal). **b**, *FBXO11* mutation types, locations, and frequencies identified in TCGA, Beat AML, TARGET pediatric AML, and AML PMP samples. **c**, *FBXO11* transcript expression in AML PMP samples with mutated/deleted (*FBXO11* mutated/loss) or WT *FBXO11* represented as a box and whisker plot. **d**, Indicated myeloid cell lines were immunoblotted for FBXO11 expression. *P* values represent two-tailed *t*-tests and error bars represent s.d.

**Supplementary Figure 3. CD34^+^ HSPC with *FBXO11* depletion have reduced cell death.**

**a**, Robust multiarray averaging (RMA)-normalized *FBXO11* signal intensity in CD34^+^ HSPC samples (*N* = 3 independent replicates). **b**, Validation of FBXO11 protein depletion by shRNA in K562 cells. **c**, The Annexin V^+^ population within total (left) or CD34^+^ (right) cells was analyzed by flow cytometry (*N* = 6 (shCTR), *N* = 7 (sh*FBXO11*) independent replicates). **d**, Representative plot of cultured CD34^+^ HSPC analyzed for expression of long-term and short-term HSC (CD90^+/-^CD45RA^-^) or hematopoietic progenitor (CD90^-^CD45RA^+^) markers. *P* values represent two-tailed *t*-tests and error bars represent s.d. **e**, Cell cycle states of the indicated HSPC populations were determined by Ki-67 and DAPI staining (*N* = 3 (shCTR), *N* = 5 (sh*FBXO11*) independent replicates).

**Supplementary Figure 4. Proteins co-immunoprecipitating with FLAG-FBXO11.**

Mean protein abundance (log_2_(ProA)) between replicates and transformed coefficient of variation (cvT) of FLAG-FBXO11 co-immunoprecipitated proteins **a**, without endonuclease (*n* = 3 independent replicates) and **b**, with endonuclease (*n* = 4 independent replicates), detected by quantitative tandem mass spectrometry. All detected proteins are plotted with the targets detected in both experimental conditions labelled.

**Supplementary Figure 5. LONP1 reciprocally co-immunoprecipitates with FBXO11.**

Co-immunoprecipitation (IP) with (**a**) FLAG IP for FLAG-FBXO11 or (**b**) LONP1 in *FBXO11*-KO K562 cells expressing empty vector (-) or FLAG-FBXO11 (+) followed by immunoblotting (IB) for LONP1 and FBXO11.

**Supplementary Figure 6. Mitochondrial mass does not decrease with *FBXO11* or *LONP1* knockdown.**

**a**, Mitochondrial mass of CD34^+^ HSPC expressing a non-targeting control shRNA (shCTR) or shRNA targeting *FBXO11* or *LONP1* was measured using detection of the MitoTracker dye by flow cytometry (*N* = 5), *P* values represent two tailed t-tests and error bars represent s.d. **b**, Validation of LONP1 protein depletion by *LONP1* targeting shRNA in K562 cells. **c**, Whole cell lysates from K562 parental cells (K562) or *FBXO11*-KO CRISPR clones expressing empty vector or FLAG-FBXO11 were immunoblotted for TOM20.

**Supplementary Figure 7. FBXO11-ΔFbox does not interact with LONP1.**

Whole cell lysates from K562 *FBXO11*-KO cells expressing empty vector, WT FLAG-*FBXO11*, or an *FBXO11* mutant lacking the F-box domain (FBXO11**-**ΔFbox) was immunoprecipitated with LONP1 or FLAG-tag antibody and the immunoprecipitated proteins or whole cell lysates (Input) were immunoblotted (IB) for FBXO11, FLAG, and αTubulin.

**Supplementary Figure 8 . Confirmation of *FBXO11* and *LONP1* transcript over-expression and/or knockdown in RNA-seq analysis.**

Transcript expression of indicated genes expressed as log_2_(CPM) (counts per million reads mapped) between (b/w) indicated experimental conditions in the CD34^+^ HSPC RNA-seq experiment

**Supplementary Figure 9. Ubiquitin pathway mutations cooccur with *RAS* and *CEBPA* mutations.**

Shown are odds ratios of AML-associated mutations for AML PMP samples occurring in ubiquitin pathway-mutated samples compared to samples that are wild-type for ubiquitin pathway genes. Mutations with odds ratio > 1 are likely to co-occur with ubiquitin pathway mutations, and odds ratios < 1 are likely to occur mutually exclusively with ubiquitin pathway mutations. *P* values determined by Fisher’s exact test, * *P*<0.05, bars represent 95% confidence intervals.

**Supplementary Figure 10. Quantification of total cell numbers and CFCs in sh *FBXO11*-, *AML1-ETO-* and *KRAS*^G12D^-transduced CD34^+^ HSPC harvested after long term culture.**

**a**, Total number of live cells harvested from long-term culture were counted using trypan blue staining. **b**, CFCs were counted on day 17 after plating (*N* = 3-12 independent replicates). *P* values represent two-tailed *t*-tests and error bars represent s.d. EV = Empty vector and control shRNA; F = shFBXO11; A = *AML1-ETO*; K = *KRAS*^G12D^; FA = F + A; KA = K + A; FKA = F + K + A. **c**, Protein abundance of FBXO11, RAS, and AML1-ETO in xenotransplanted combinations.

**Supplementary Figure 11. Analysis of bone marrow and spleen cells from primary xenotransplanted mice at endpoint.**

**a**, Percentages of double CD45^+^ human hematopoietic cells using two distinct antibody clones in bone marrow at endpoint. **b**, Spleen weights were measured at endpoint (*n* = 9-13 mice). **c**, Representative spleens are shown. **d**, Total live bone marrow cell counts at endpoint (*n* = 9-13 mice). *P* values represent two-tailed *t-*tests. Error bars represent s.d.

**Supplementary Figure 12. Myeloid cell infiltration in the leptomeninges of FKA mice.**

Myeloperoxidase stains performed on brain sections of a wild-type NRG-3GS mouse (Control) and an FKA mouse, showing myeloid cell infiltration.

**Supplementary Figure 13. Flow cytometry analysis of bone marrow cells harvested at endpoint.**

**a**, Flow cytometry analysis of lymphoid and myeloid cell engraftment at endpoint following xenotransplantation (*n* = 5 mice). *P* values represent two-tailed *t-*tests and error bars represent s.d. **b**, Percentages of CD34^+^ cells within the engrafted human hematopoietic cell population in bone marrow at endpoint (*n* = 5-12 mice, ND = not detected. *P* values represent Fisher’s exact test, error bars represent s.d.).

**Table S1.** Additional oncoprint mutation information, related to Figure 1.

**Table S2**. FBXO11 mutations in non-AML TCGA cancers, related to Figure S2.

**Table S3**. SCF gene expression volcano plot, related to figure Figure 2.

**Table S4**. GSEA for pathways enriched with FBXO11 depletion in CD34^+^ HSPC, related to Figure 3.

**Table S5**. Co-immunoprecipitating proteins with FLAG-FBXO11 detected by MS/MS, related to Figure 4.

**Table S6**. CD34^+^ cell quantification from LTC to xenotransplant endpoint, related to Figure 7.

---

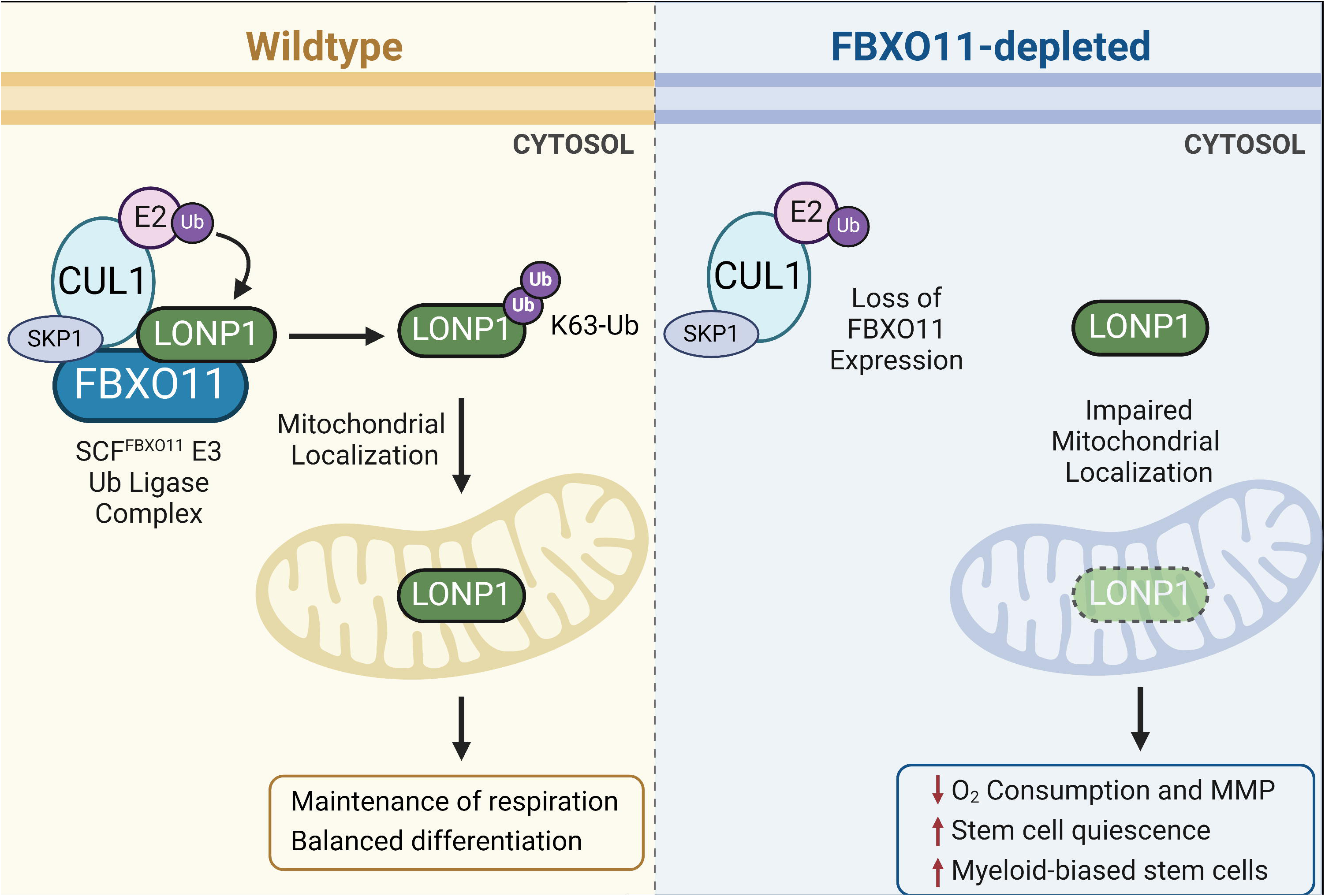

## Notes

### Competing Interest Statement

The authors have declared no competing interest.

### Summary of Updates

We have substantially updated the manuscript to reflect its current state, including removing a main figure and adding two new figures.

