## Supplementary material for "Loss of FBXO11 function establishes a stem cell program in acute myeloid leukemia through dysregulation of mitochondrial LONP1": Figures S1-S13

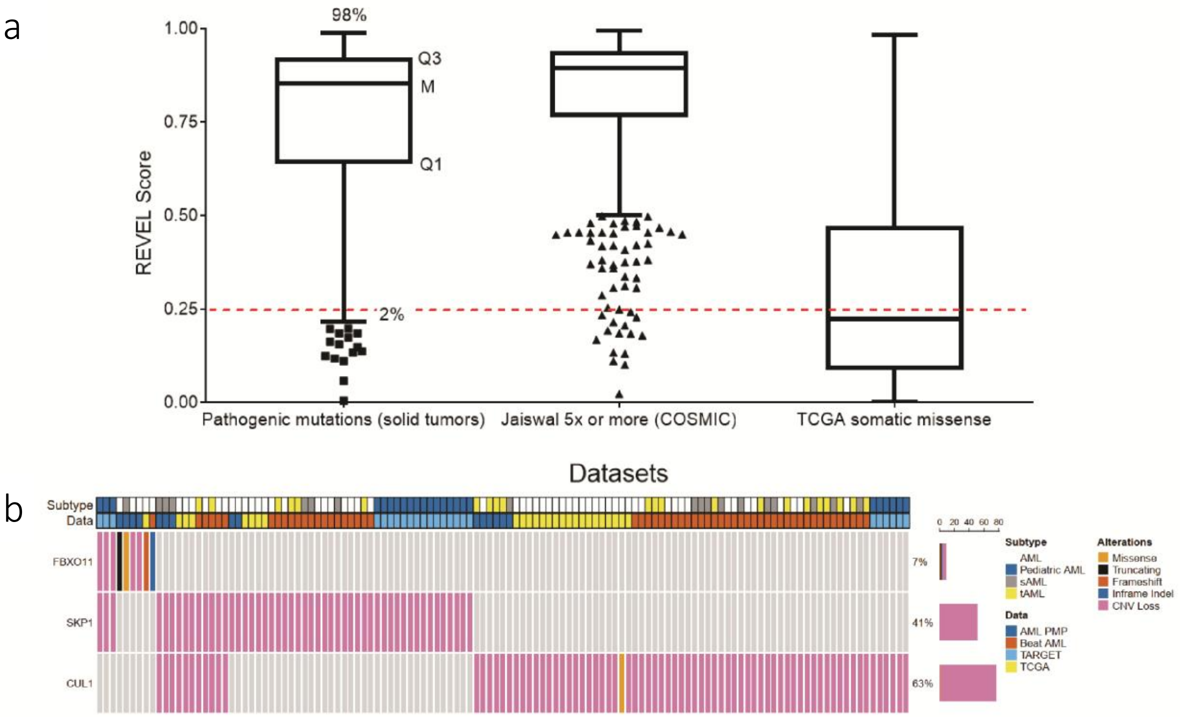


**Supplementary Figure 1. Mutations affecting the ubiquitin pathway occur frequently in AML.**

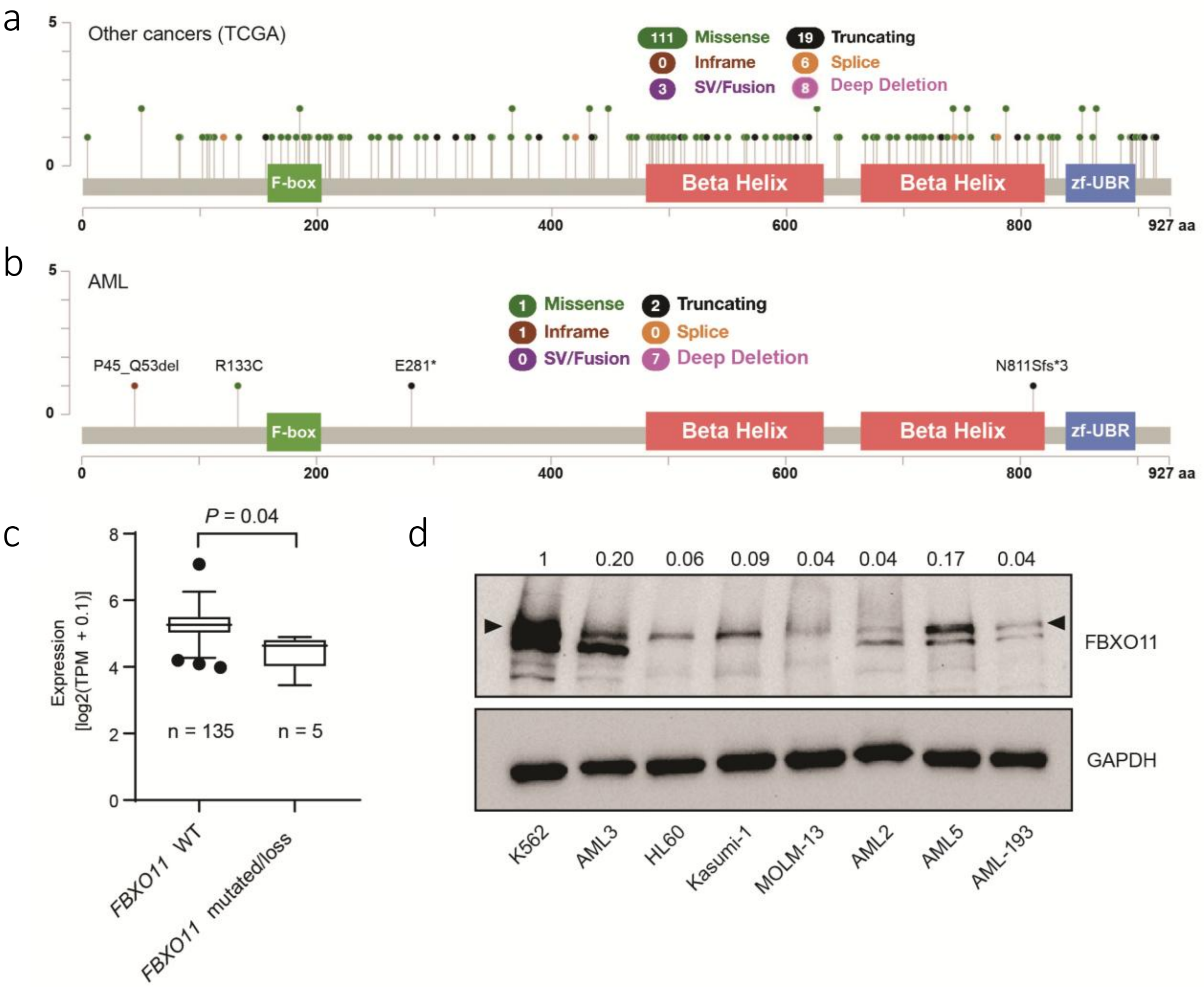


**Supplementary Figure 2.** ***FBXO11*** **is recurrently mutated in multiple cancers.**

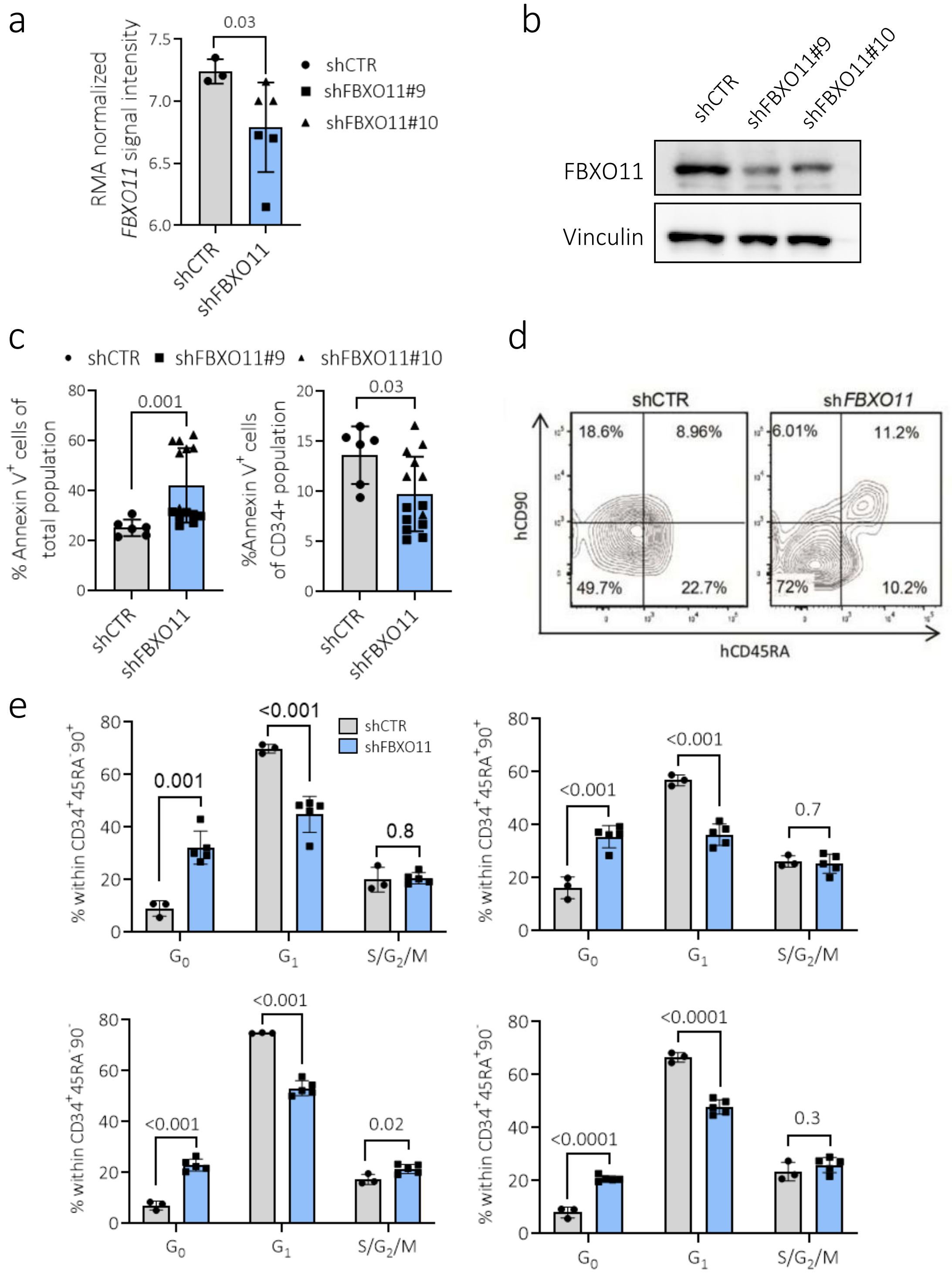


**Supplementary Figure 3. CD34+ HSPC with *FBXO11* depletion have reduced cell death.**

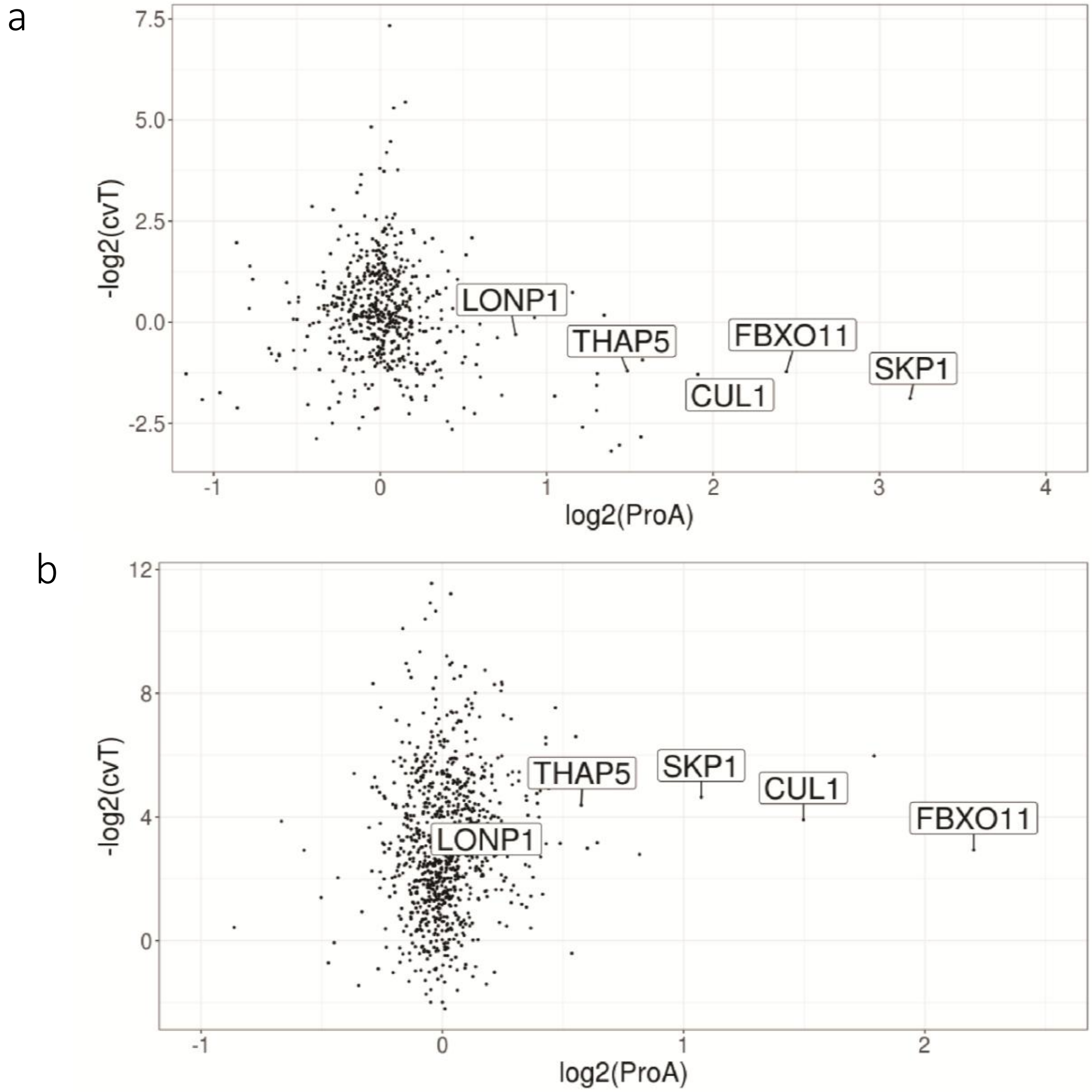


**Supplementary Figure 4. Proteins co-immunoprecipitating with FLAG-FBXO11.**

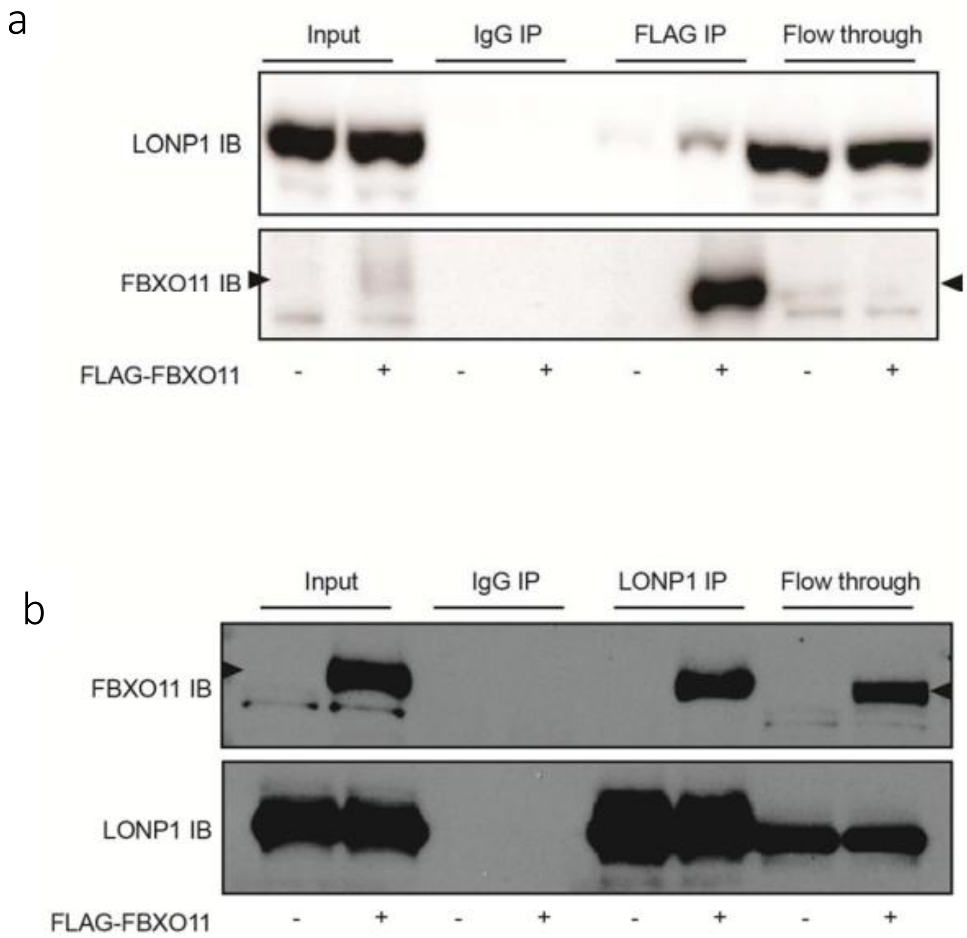


**Supplementary Figure 5**. **LONP1 reciprocally co-immunoprecipitates with FBXO11.**

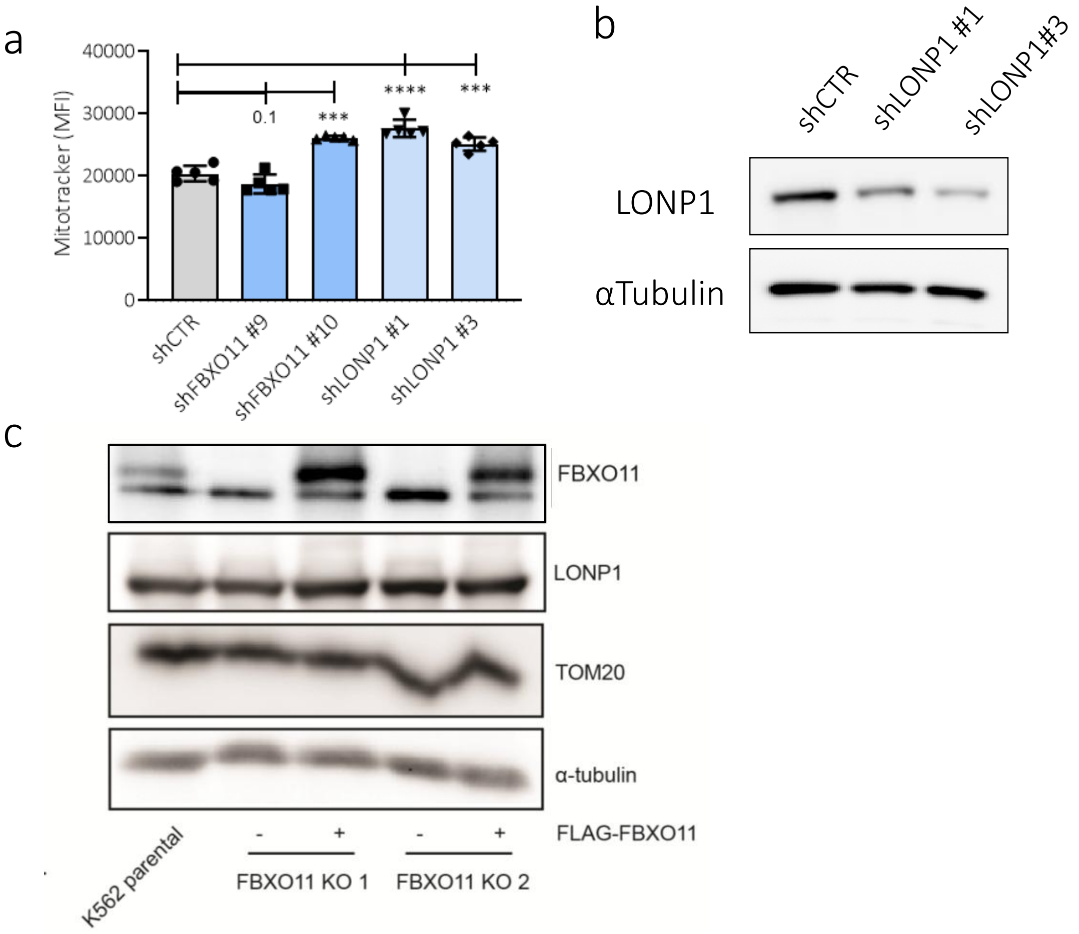


**Supplementary Figure 6**. **Mitochondrial mass does not decrease with *FBXO11* or *LONP1* knockdown.**

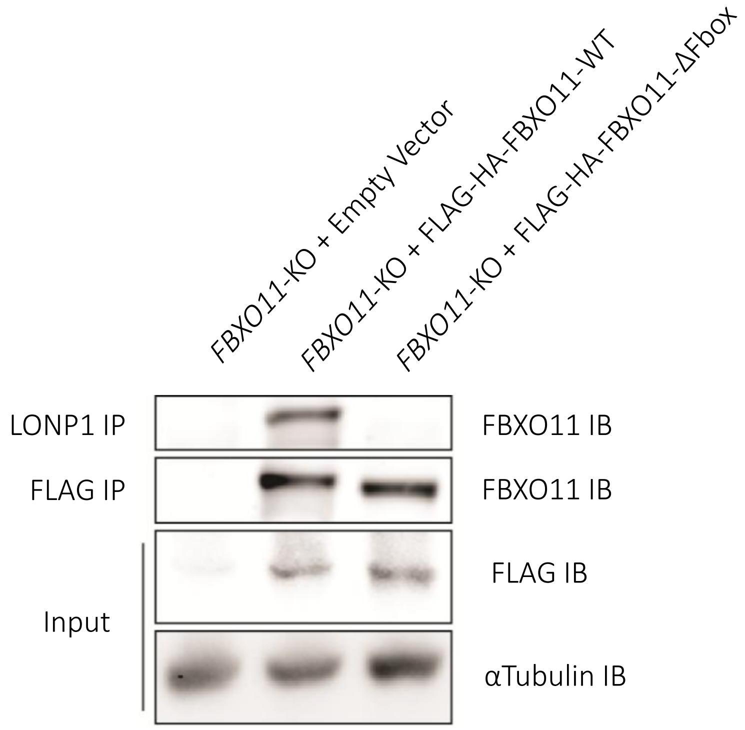


**Supplementary Figure 7. FBXO11-ΔFbox does not interact with LONP1.**

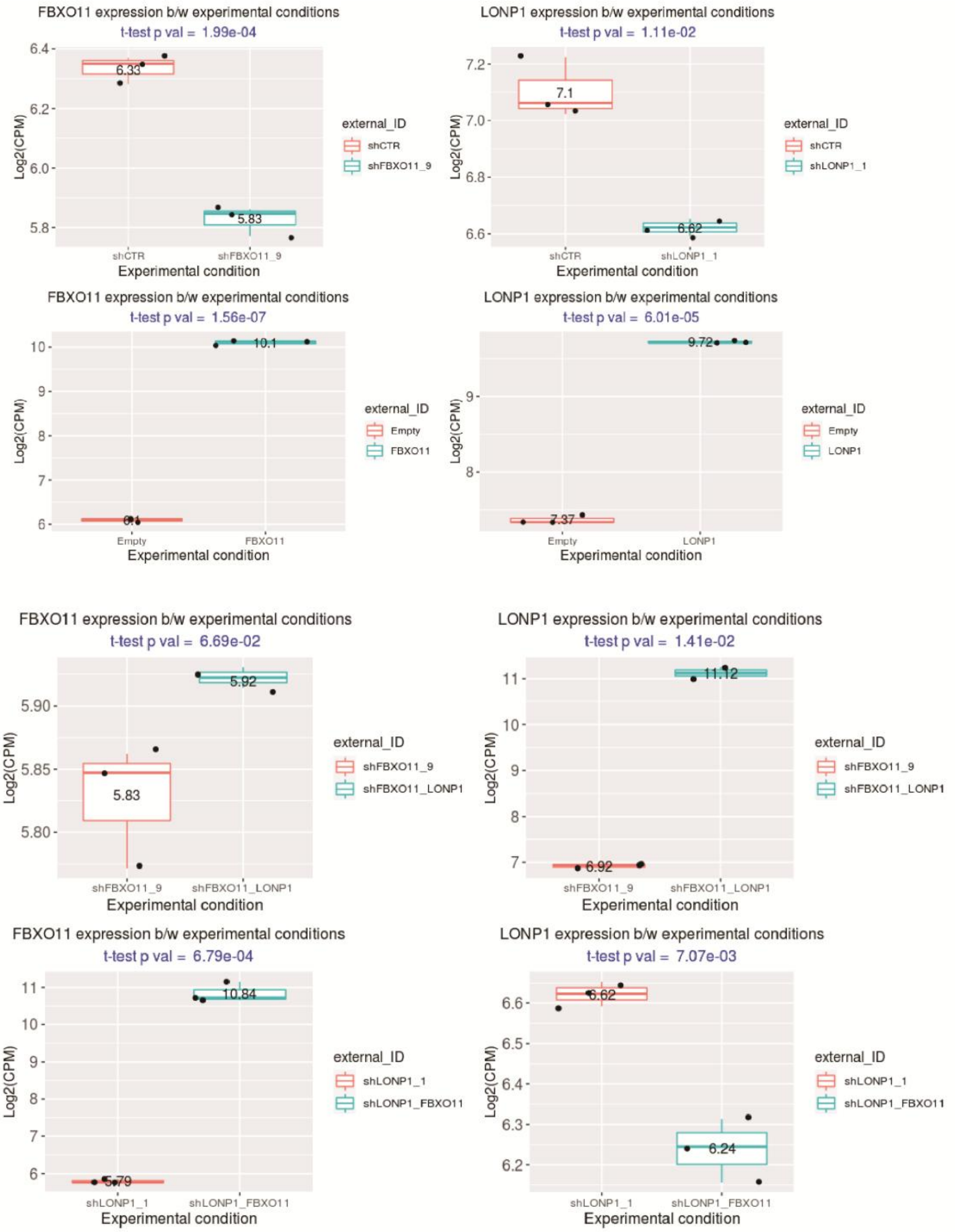


**Supplementary Figure 8**. **Confirmation of *FBXO11* and *LONP1* transcript over-expression and/or knockdown in RNA-seq analysis.**

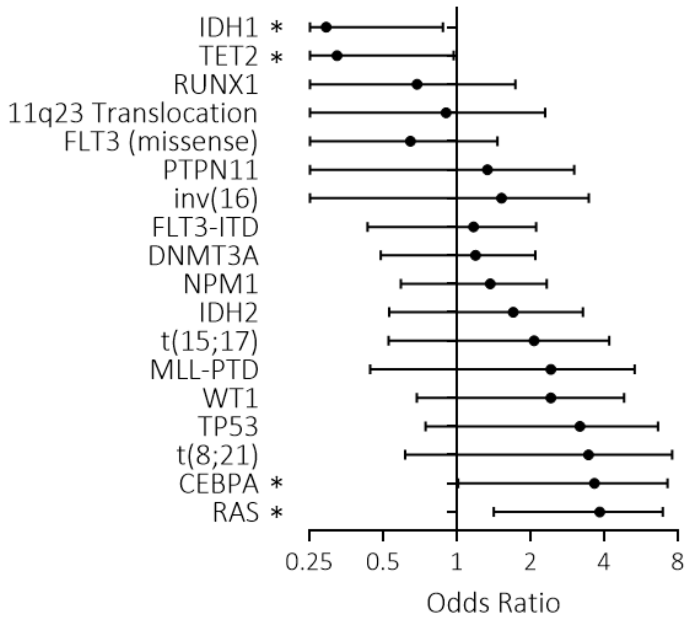


**Supplementary Figure 9**. **Ubiquitin pathway mutations cooccur with *RAS* and *CEBPA* mutations.**

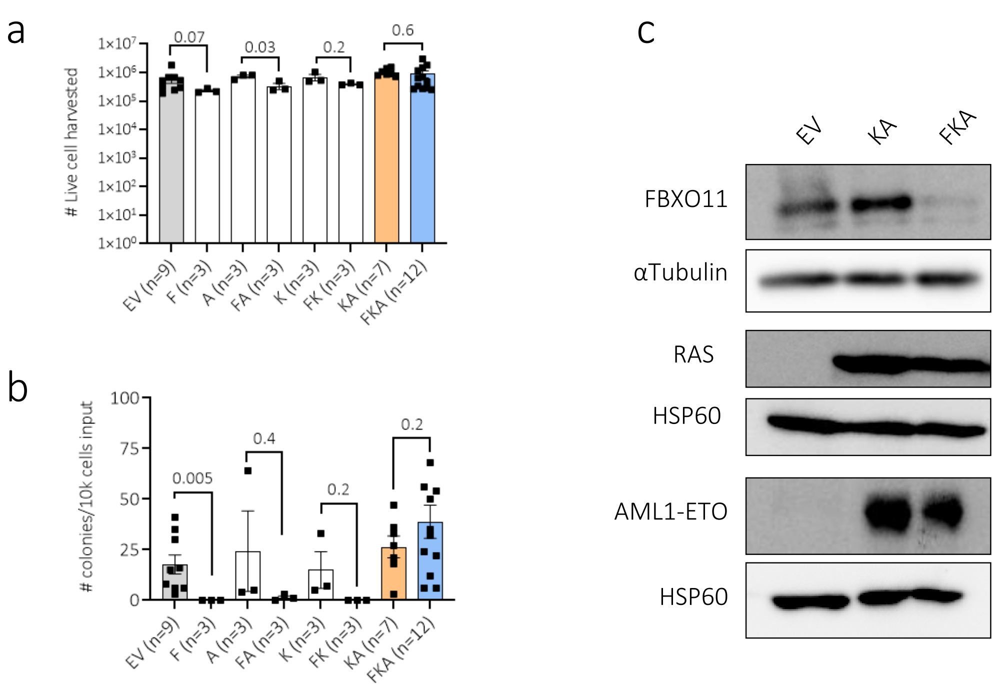


**Supplementary Figure 10. Quantification of total cell numbers and CFCs in sh*FBXO11*-, *AML1-ETO*- and *KRAS*$^{G12D}$- transduced CD34$^{+}$ HSPC harvested after long term culture.**

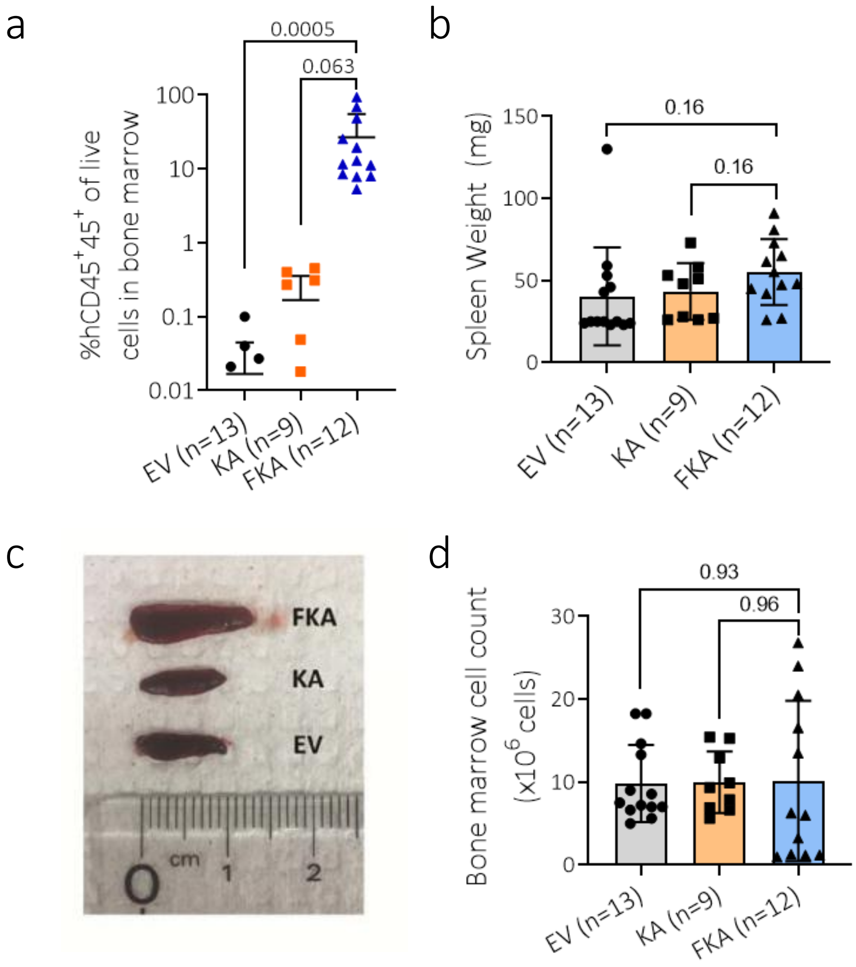


**Supplementary Figure 11. Analysis of bone marrow and spleen cells from primary xenotransplanted mice at endpoint.**

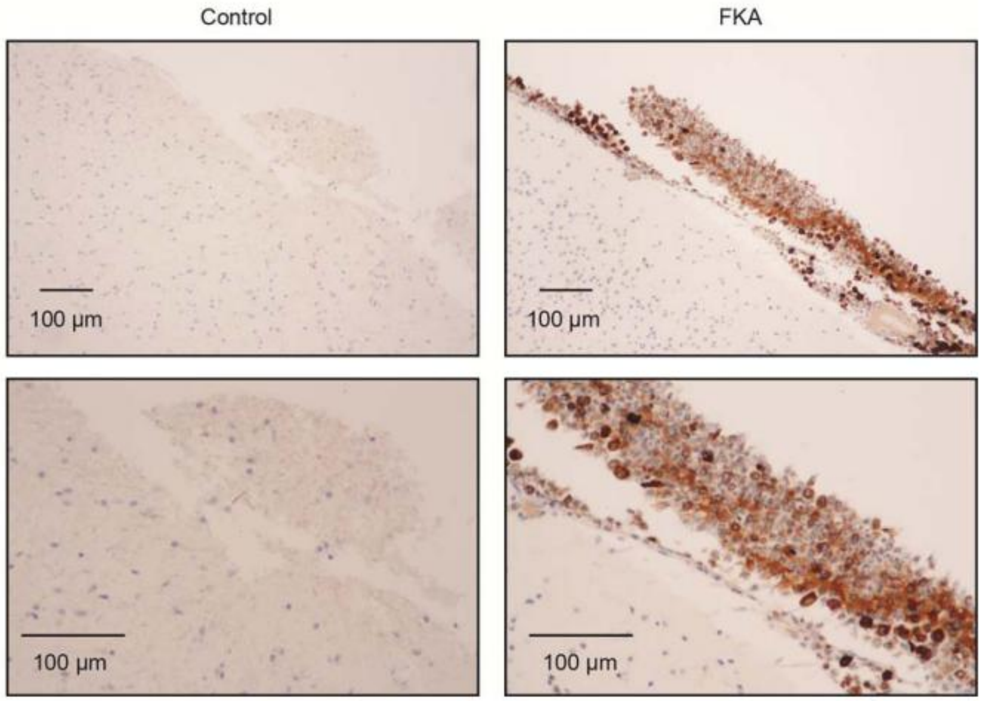

**Supplementary Figure 12. Myeloid cell infiltration in the leptomeninges of FKA mice.**

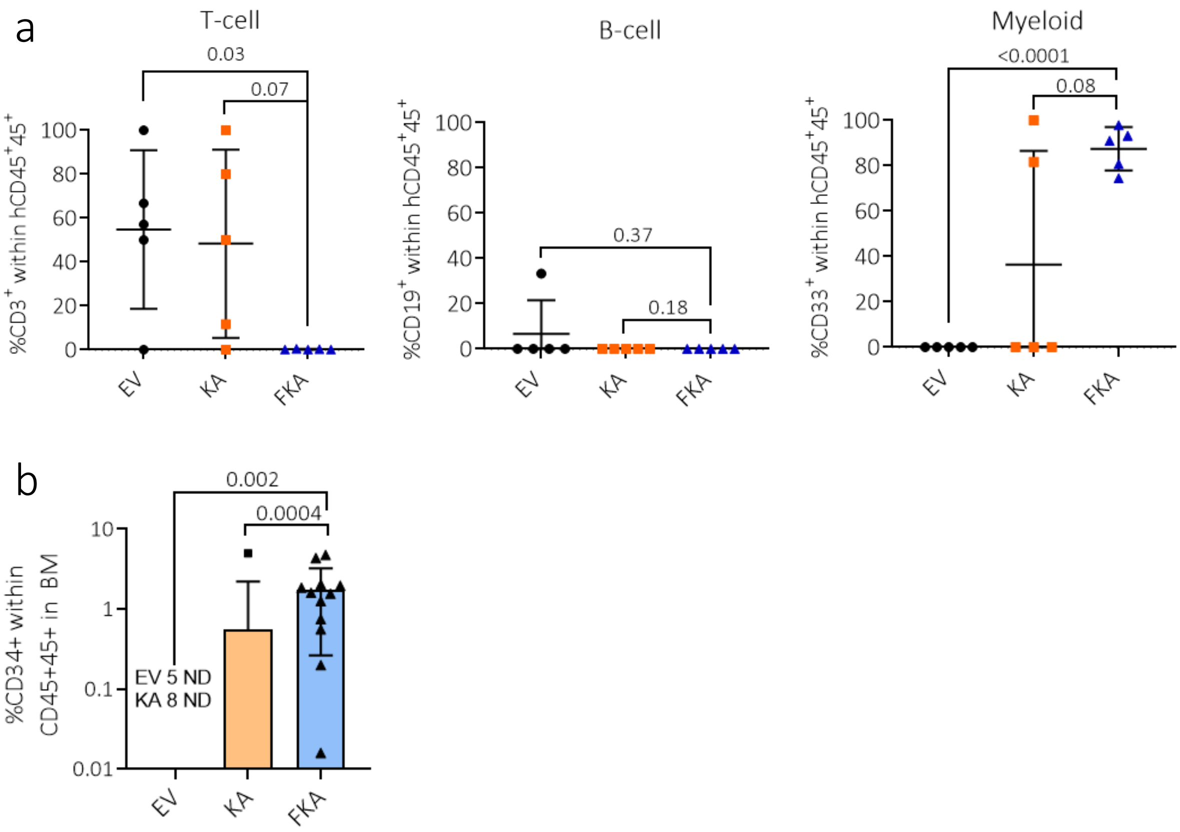


**Supplementary Figure 13. Flow cytometry analysis of bone marrow cells harvested at endpoint.**
